# The extracellular heparan sulfatase SULF2 limits myeloid IFNβ signaling and Th17 responses in inflammatory arthritis

**DOI:** 10.1101/2024.03.13.584823

**Authors:** Maarten Swart, Andia Redpath, Joy Ogbechi, Ryan Cardenas, Louise Topping, Ewoud B. Compeer, Michael Goddard, Anastasios Chanalaris, Richard Williams, Daniel S. Brewer, Nicola Smart, Claudia Monaco, Linda Troeberg

**Affiliations:** Kennedy Institute of Rheumatology, Nuffield Department of Orthopaedics, Rheumatology and Musculoskeletal Sciences, University of Oxford, Roosevelt Drive, Headington, Oxford, OX3 7FY, UK; Department of Physiology, Anatomy & Genetics, University of Oxford, Sherrington Building, Oxford, OX1 3PT, UK; Norwich Medical School, University of East Anglia, Bob Champion Research and Education Building, Rosalind Franklin Road, Norwich, NR4 7UQ, UK

**Keywords:** heparan sulfate, interferon, inflammation, macrophage

## Abstract

Heparan sulfate (HS) proteoglycans are important regulators of cellular responses to soluble mediators such as chemokines, cytokines and growth factors. We profiled changes in expression of genes encoding HS core proteins, biosynthesis enzymes and modifiers during macrophage polarisation, and found that the most highly regulated gene was *Sulf2*, an extracellular HS 6-O-sulfatase that was markedly downregulated in response to pro-inflammatory stimuli. We then generated *Sulf2*^+/-^ bone marrow chimeric mice and examined inflammatory responses in antigen-induced arthritis, as a model of rheumatoid arthritis. Resolution of inflammation was impaired in myeloid *Sulf2*^+/-^ chimeras, with elevated joint swelling and increased abundance of pro-arthritic Th17 cells in synovial tissue. Transcriptomic and *in vitro* analyses indicated that *Sulf2* deficiency increased type I interferon signaling in bone marrow-derived macrophages, leading to elevated expression of the Th17-inducing cytokine IL-6. This establishes that dynamic remodeling of HS by *Sulf2* limits type I interferon signaling in macrophages, and so protects against Th17-driven pathology.

## Introduction

Inflammation is regulated by a network of cytokines, chemokines and growth factors that control the survival, proliferation, recruitment and signaling responses of innate and adaptive immune cells. Many of these extracellular mediators bind to heparan sulfate (HS) proteoglycans [1], and this interaction controls their localisation, stability, gradient formation and/or signaling [2–4]. For example, binding to extracellular matrix HS is required for the *in vivo* chemotactic activity of CCL2, CCL4 and CCL5 [5], and regulates the *in vivo* half-life [6] and bioactivity of IFNγ [7]. While less well-studied, cell surface HS has also been shown to modulate the responses of innate and adaptive immune cells to cytokines, chemokines and growth factors. For example, B cells require cell surface HS to proliferate and differentiate in response to A Proliferation-Inducing Ligand (APRIL) [8, 9].

HS is a long, structurally diverse polysaccharide that can bind to >400 protein ligands [1] with a range of affinities, and consequently a broad range of biological effects [10, 11]. Unlike the template-dependent synthesis of nucleic acids and proteins, synthesis of heparan sulfate is a template-independent process, in which up to 20 enzymes act sequentially in the Golgi apparatus to add alternating uronic acid (either glucuronic acid or iduronic acid) and N-acetylglucosamine residues to the growing HS chain and sulfate it at various positions [11] i.e. glucuronic acid can be 2-O-sulfated (by HS2ST1), while N-acetylglucosamine can be N-sulfated (by NDST1-4), 6-O-sulfated (by HS6ST1-3) or 3-O-sulfated (by HS3ST1-6) during HS biosynthesis [11]. After it has been synthesised and secreted from cells, HS can also be modified in the extracellular environment by heparanase, an enzyme that degrades the HS chains [11], or by the sulfatases SULF1 and SULF2, which specifically remove sulfate groups from the 6-carbon position of N-acetylglucosamine residues [11]. Changes in expression of these biosynthetic and modifying enzymes lead to dynamic changes in HS structure and function during multiple physiological (development [12], wound healing and ageing [13–15]) and pathological (diabetes [16], fibrosis [17, 18], Alzheimer’s disease [19], osteoarthritis [20], cancer [18, 21]) processes.

In myeloid cells, expression of HS sulfotransferases has been shown to change in response to cytokine-induced polarisation [22–24], leading to changes in HS abundance and sulfation [23, 25]. A key study by Gordts *et al*. [26] showed that myeloid-specific deletion of *Ndst1* sensitised macrophages to type I interferon (IFN) signaling, and exacerbated inflammation in murine models of atherosclerosis and obesity. *In vitro* mechanistic studies indicated that *Ndst1*-dependent N-sulfation increased IFNβ binding to HS, potentially sequestering the cytokine from its cognate IFN receptors [26]. Subsequent studies showed *Ndst1* can also quench inflammatory responses in dendritic cells (DCs), with deletion of *Ndst1* in DCs increasing antigen presentation and cytotoxic T cell responses in a lung carcinoma model [27].

Here, we systematically profiled changes in expression of the 12 HS core proteins and 25 HS biosynthesis and modifying enzymes in murine bone marrow-derived macrophages (BMDMs) in response to pro- and anti-inflammatory stimuli, and found that the 6-O-sulfatase *Sulf2* was the most highly regulated gene. Using a bone marrow transplantation strategy, we generated chimeric mice with myeloid *Sulf2*-deficiency and examined inflammatory responses *in vivo* in an antigen-induced arthritis model. This showed that myeloid SULF2 limits inflammation, by suppressing type I interferon signaling and so reducing arthritogenic Th17 responses.

## Materials and methods

### Reagents

IFNγ (BioLegend), IL1, IL2, IL4, IL6, IL13, IL23, macrophage colony-stimulating factor (M-CSF), granulocyte-macrophage colony-stimulating factor (GM-CSF) and transforming growth factor beta (TGFβ) were all purchased from Peprotech, and ultra-pure lipopolysaccharide (LPS) from *Escherichia coli* O111:B4 was purchased from InvivoGen RT-qPCR primers are detailed in Supplementary Tables S1 and S2, with antibodies used for flow cytometry shown in Supplementary Table S3.

### Primary cell isolation and culture

All experiments were carried out in accordance with the United Kingdom Home Office Animals Scientific Procedures Act 1986 under relevant personal and project licenses.

Single cell suspensions were generated from bone marrow by flushing femurs and tibias with Roswell Park Memorial Institute 1640 medium (RPMI), and adding red blood cell lysis buffer (2 min, room temperature, Sigma). Cells were cultured (7 d) in complete medium [RPMI supplemented with 10 % fetal bovine serum (FBS), 1 % penicillin/streptomycin, 50 μM 2-mercaptoethanol] supplemented with M-CSF (100 ng/ml, Peprotech) to generate BMDM), or with GM-CSF (20 ng/ml, Peprotech) to generate dendritic cells (DCs). BMDMs were replated on day 7, and stimulated for 18 h with IL-4 (20 ng/ml, Peprotech) plus IL-13 (20 ng/ml, Peprotech), or with LPS (100 ng/ml, InvivoGen) plus IFNγ (100 ng/ml, BioLegend).

Single cell suspensions were generated from joint synovium by digesting opened hindleg knee joints in liberase and DNase (30 min, 37 °C), and from lungs by digestion with collagenase IV and DNase I (40 min, 37 °C). Single cell suspensions were generated from spleen and lymph nodes by pressing tissues through a 70 μm cell strainer. All single cell suspensions, except those use for generating chimeras, were treated with red blood cell lysis buffer (2 min, room temperature, Sigma) before further use.

### Generation of *Sulf2*^+/-^ mice

*Sulf2*^+/-^ mice were generated from the *Sulf2*^tm1a(KOMP)Wtsi^ embryonic stem cell line (RRID:MMRRC_063144-UCD, obtained from the KOMP Repository at University of California, Davis), in which exon 4 of the *Sulf2* gene is flanked by LoxP sites. Embryonic stem cells were injected into blastocysts and resulting chimeras bred to germline transmission of the *Sulf2* tm1a allele, before crossing with Tg(Pgk1-cre)Lni mice [63], available from JAX, to facilitate recombination, and backcrossing to a C57BL/6 background to generate *Sulf2* tm1b allele mice, with a reporter-tagged deletion allele, and missing critical exon 4 of the *Sulf2* gene. The deletion of exon 4 is expected to generate a mis-sense mutation and to cause a premature stop codon in exon 5, which is anticipated to lead to nonsense-mediated decay of the mRNA and loss of protein expression.

### RT-qPCR

RNA was reverse transcribed using a High Capacity Reverse Transcription cDNA kit (Applied Biosystems) after extraction from cultured cells (with RNeasy mini spin-columns, Qiagen), tissues (with TRIzol, Life Technologies), or joints [by homogenisation in a TissueLyser II using a Precellys hard tissue Lysing Kit (Bertin) followed by RNeasy mini spin-columns, Qiagen]. TaqMan Low-Density Array (TLDA) cards (Applied Biosystems) were custom designed using TaqMan primer/probes (Applied Biosystems), with data acquired on a ViiA Real-Time PCR System (Applied Biosystems). Manual RT-qPCR was carried out using TaqMan primer/probes (Applied Biosystems) with data acquired on a ViiA Real-Time PCR System (Applied Biosystems, Supplementary Table S1), or with KiCqStart SYBR Green Primers with data acquired and melt curves examined on a QuantStudio 3 (ThermoFisher Scientific, Supplementary Table S2). Fold change in expression was calculated using the ΔΔCt method.

### Flow cytometry

Cells were stained with Zombie near-infra red (BioLegend) or aqua (ThermoFisher Scientific) fixable live/dead stains (10 min, room temperature), washed in PBS and blocked with anti-mouse CD16/CD32 (10 min, 4 °C, BD Biosciences). For extracellular staining, cells were then incubated with fluorochrome-conjugated antibodies (Supplementary Table S3) in PBS with 1 % FBS (4 °C, 30 min), washed twice, and fixed in BD CellFIX (10 min, on ice, BD Biosciences). For intracellular staining, cells were fixed in Foxp3/Transcription factor staining buffer (4 °C, 30 min, Invitrogen), and incubated with fluorochrome-conjugated antibodies in Permeabilisation buffer (4 °C, 30 min, Invitrogen). To quantify intracellular cytokines, cells were stimulated (4 h) with phorbol 12-myristate 13-acetate (PMA, 20 ng/ml) plus ionomycin (1 mg/ml) in the presence of GolgiPlug/GolgiStop (BD Biosciences). Fluorescence was quantified using an LSR-II or Fortessa X20 flow cytometer (BD Biosciences).

### Phagocytosis assays

BMDMs were incubated with 1 μm fluorescent microspheres [Fluoresbrite plain yellow-green particles, multiplicity of infection (MOI) 50, 30 min, Polysciences Inc], fluorescein-conjugated *Escherichia coli* K-12 BioParticles (MOI 5, 15 min, Invitrogen), or fluorescein-conjugated *S. cerevisiae* zymosan A Bioparticles (MOI 5, 20 min, Invitrogen), and phagocytosis quantified by flow cytometry. Efferocytosis was quantified by incubating BMDMs with Jurkat T cells (MOI 2, 45 min) which had been labelled with calcein-acetoxymethyl (2 h, 37 °C, ThermoFisher Scientific) and exposed to UV light (305 nm, 2.5 h) to induce apoptosis. As a negative control for all phagocytosis assays, particles were also incubated with BMDMs on ice.

### Generation of bone marrow chimeras and antigen-induced arthritis (AIA)

Recipient 8-week old wild-type (WT) C57BL/6 mice were irradiated (two doses of 5.5 Gy at a 4 h interval in a Gulmay X-ray irradiator), and injected 2 h later in the tail vein with bone marrow cells (5 million/mouse) from 8-week old WT or *Sulf2*^+/-^ donors. Seven weeks later, mice were subcutaneously injected with methylated bovine serum albumin (mBSA, 100 μg, Sigma) emulsified 1:1 with Complete Freund’s adjuvant (BD Biosciences). Arthritis was induced 3 weeks later by intra-articular tibiofemoral injection of mBSA (100 μg in PBS, right knee) or PBS control (left knees). Knee swelling was measured daily for 7 days after intra-articular injection using a calliper, and pain assessed by measuring weight bearing on mBSA-injected compared to PBS-injected paws using a dynamic weight-bearing 2.0 chamber (Bioseb). Mice were sacrificed after 7 days, and immune infiltration and total histological score, composed of the sub-synovium, synovium and bone marrow density scores, calculated [64]. To analyse T cell subsets, single cell suspensions from knee joints and lymph nodes were stimulated for 4 h with PMA (20 ng/ml) and ionomycin (1 mg/ml) in the presence of protein transport inhibitors, before CD3+, CD4+ and CD8+ subsets were analysed by flow cytometry.

### Bulk RNASeq

RNA was isolated (RNeasy Micro Kits, Qiagen) from single cell suspensions prepared from knee joints of WT and *Sulf2*-deficient bone marrow chimera mice sacrificed 7 days after induction of arthritis. Oxford Genomics Centre prepared bulk RNA libraries using an NEBNext Single Cell/Low Input RNA Library Prep Kit for Illumina (New England Biolabs) and sequenced the samples at a depth of 25 million reads per sample on a NovaSeq6000 (Illumina). PolyA transcripts were analysed with the Nextflow RNA-Seq pipeline [65], using STAR for alignment with the GRCm38 reference murine genome. Deconvolution of cell types was performed with MuSIC [32], utilising single cell data from Park *et al.* [66]. DeSeq2 was used for differential gene expression analysis, with correction for multiple comparisons carried out using the Benjamini and Hochberg method. Functional enrichment analysis was performed using gProfiler2 (v0.2.0) utilising the Kyoto Encyclopedia of Genes and Genomes (KEGG) pathway database [67] for terms. The gSCS (Set Counts and Sizes) correction method was used to determine significantly enriched pathways with significance p<0.05. Differentially regulated genes were manually compared with the Interferome database interferome.org, [33]), and Homer was used to analyse promoter regions (8 base pair motifs, −400 to −100 base pairs upstream of transcriptional start sites) of differentially expressed genes [68].

### Analysis of type I interferon signaling

For analysis of STAT1 phosphorylation, BMDM from WT and *Sulf2^+/-^* mice were stimulated with murine IFNβ (50 ng/ml, 30 min, Peprotech) and phospho-STAT1 (Y701) (D4A7, Cell Signaling) and STAT1 (#9172, Cell Signaling) quantified by immunoblotting analysis on an Odyssey CLx fluorescent imaging system (LI-COR Biosciences) using Image Studio software (ver 4.0, LI-COR Biosciences).

For analysis of interferon-regulated gene expression, BMDM from WT and *Sulf2^+/-^* mice were stimulated with murine IFNβ (50 ng/ml, 4 h, Peprotech) and expression of *Ccl5*, *Ccl7*, *Tlr3* and *Il6* quantified by RT-qPCR relative to *Gapdh*.

### Primeflow analysis of *Sulf2* mRNA expression in blood, bone marrow, spleen, and lung

The PrimeFlow RNA assay (Invitrogen) was used according to the manufacturer’s protocol to assess *Sulf2* mRNA expression in various cell types. Isolated cells were surface-stained, permeabilised, fixed, and hybridised with a target probe set (2 h, 40 °C). After hybridisation of pre-amplifying primers (1.5 h, 40 °C) and amplifying primers (1.5 h, 40 °C), samples were incubated with labelled probes (1 h, 40 °C), stained for immune cell markers (CD45, CD3, CD19, Ly6C, Ly6G in blood and bone marrow; CD45, CD3, CD19, F4/80, CD11b, Ly6C, Ly6G, CD11c, CD64, MHC-II in spleen and lung) and fluorescence quantified using a Fortessa X20 flow cytometer (BD Biosciences).

### Toll-like receptor (TLR) signaling

Macrophages were stimulated with the TLR agonists IFNγ, LPS, IFNγ+LPS (all at 100 ng/ml), fibroblast-stimulating lipopeptide-1 (FSL-1, 100 ng/ml), 10 ng/ml polyinosinic:polycytidylic acid [poly (I:C)], or 100 ng/ml N-palmitoyl-S-[2,3-bis(palmitoyloxy)-(2RS)-propyl]-[R]-cysteinyl-[S]-seryl-[S]-lysyl-[S]-lysyl-[S]-lysyl-[S]-lysine (Pam3CSK4) for 6 h and IL6 secretion measured by ELISA (R&D Systems).

### Inflammasome activation

Macrophages were stimulated with 100 ng/ml LPS (100 ng/ml, 6 h) with nigericin (5 μM) added for the last 2 h of culture. Cell death was assessed using the CytoTox 96 Cytotoxicity assay (Promega) and IL1β secretion measured by ELISA (Invitrogen).

### Autophagy

Macrophages were stimulated with LPS (100 ng/ml, 18 h) and then treated with bafilomycin (100 nM, 2 h) before microtubule-associated protein light chain 3 (LC3)-I and LC3-II levels in cell lysates was analysed by immunoblotting (rabbit polyclonal LC3 antibody, Sigma).

### Antigen uptake and presentation

Antigen uptake was measured by flow cytometry analysis of BMDMs incubated with Alexa647-labelled ovalbumin (OVA, 0.1 mg/ml, 1 h, 37 °C, Molecular Probes). To assess antigen presentation, CD4+ T-cells were enriched from the spleen of OT-II mice using the MojoSort mouse CD4+ T-cell isolation kit (BioLegend) and labelled with CellTrace Violet (5 μM, 20 min, 37 °C, Invitrogen). Labelled CD4+ T-cells were co-cultured (4 d) with WT or *Sulf2*^+/-^ antigen-presenting cells (BMDM or DCs) activated with LPS (100 ng/ml LPS, 4 h, InvivoGen). As a source of antigen, OVA (0.1 mg/ml, InvivoGen) was added to antigen-presenting cells (1 h, 37 °C) before LPS-activation, or OVA peptide 323-339 (10 μg/ml, Peptides International) was added during co-culture of antigen-presenting cells with CD4+ T-cells. CD4+ T-cell proliferation and expression of CD25 were analysed by flow cytometry, with division index, proliferation index and percent divided cells determined using the FloJo proliferation tool.

### Cytotoxicity assay

A CytoTox 96 Cytotoxicity assay (Promega) was performed according to manufacturer’s instructions. This assay detects release of lactate dehydrogenase from damaged cells.

### Statistical analysis

Generation of heatmaps and principal component analysis (PCA) analysis was performed in R (version 3.5.2) and R studio (version 1.1.453). For PCA analysis, the prcomp function was used, without any maximum number of ranks and without clustering. The samples were then plotted on the obtained and rotated principal components, together with ellipsoids indicating 95% confidence around the centroids of the data groups. All other data were analysed using GraphPad Prism (version 8.4.1). Data were analysed for normality using the D’Agostino-Pearson test, and for statistical significance using relevant tests as detailed in the figure legends. Data are presented as mean ± SEM or median ± IQR where appropriate.

## Results

### Macrophage expression of HS core proteins and modifying enzymes was regulated by polarisation *in vitro*

We profiled the expression of genes encoding the 37 HS core proteins and modifying enzymes in BMDM, and found that they expressed detectable levels of 22 of these genes (8 of the possible 12 core proteins and 14 of the possible 25 modifying enzymes). Polarisation of the cells *in vitro* with IFNγ+LPS or IL4+IL13 significantly altered expression of 60% of the expressed genes (Fig. 1A), with 3 distinct groups on a PCA analysis (Fig. 1B).

**Fig. 1.**
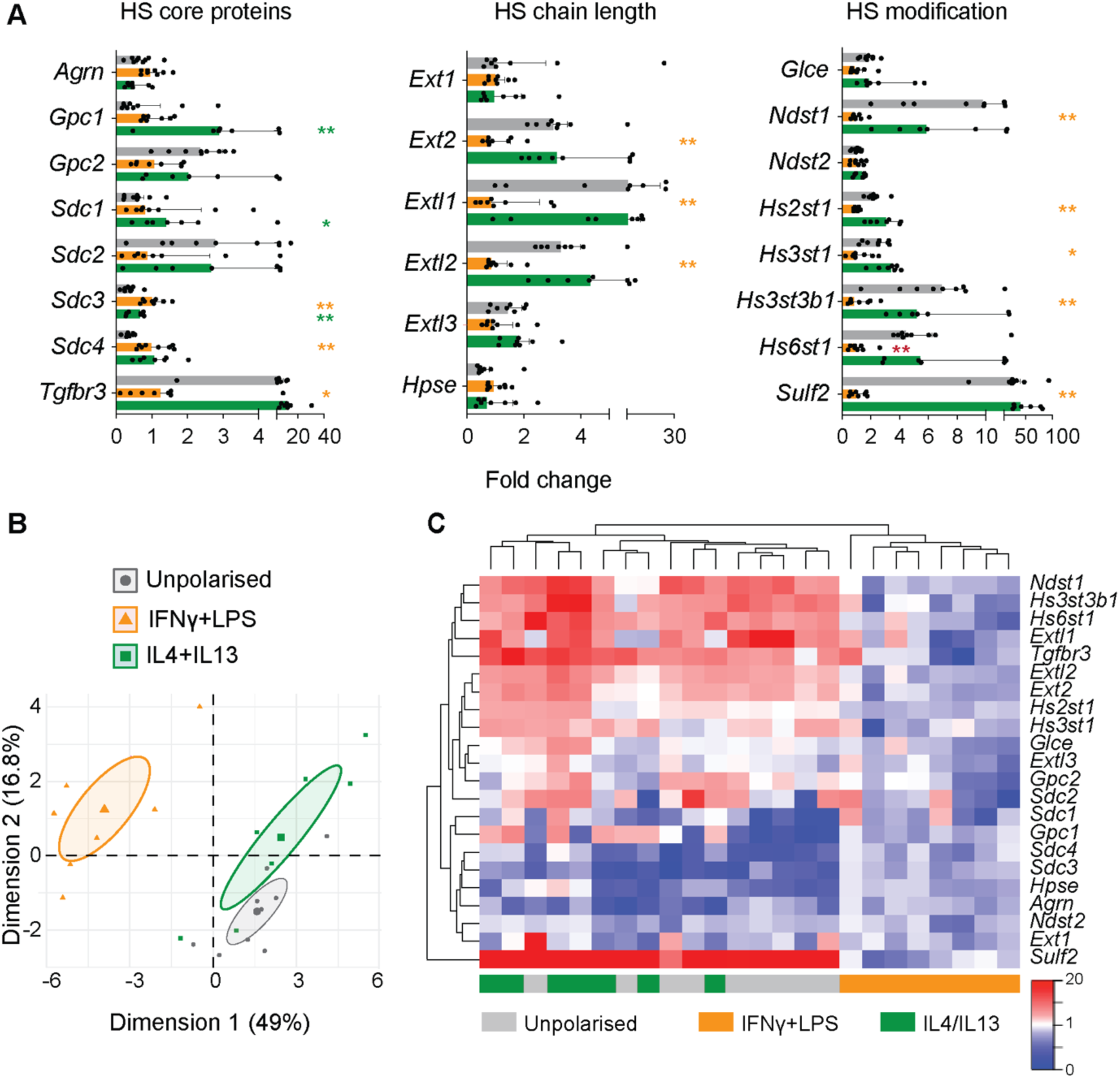
Expression of HS core proteins and modifying enzymes was downregulated by IFNγ+LPS. (**A-C**). Bone marrow-derived macrophages were polarised *in vitro* with IFNγ+LPS or IL4+IL13 (18 h, n=7-9 mice per group). mRNA expression of the 37 HS core proteins and modifying enzymes was quantified by RT-qPCR using custom-designed TaqMan low-density array cards, and expressed relative to the geometric mean of the housekeeping genes *18s*, *Gapdh*, and *Rplp0*, and to the average expression in IFNγ+LPS-polarised cells. (**A**). Fold change in expression of the 22 detectably-expressed HS core proteins and modifying enzymes (median ± IQR, analysed by Kruskall-Wallis test followed by multiplicity correction using the two-stage step-up method of Benjamini, Krieger and Yekutieli. *, P<0.05; **, P<0.01, shown in orange for IFNγ+LPS and in green for and IL4+IL13). (**B**). PCA analysis of differentially expressed genes, with ellipses indicating the 95% confidence interval. (**C**). Heatmap showing changes in expression of the 22 detectably-expressed HS genes.

Polarisation with IFNγ+LPS generated a distinct gene expression profile (Fig. 1B-C), with significantly altered expression of three core proteins (increased *Sdc3* and *Sdc4*, and reduced *Tgfbr3* compared to unpolarised BMDM) and lower expression of many of the HS modifying enzymes (Fig. 1A). At least one gene from each of the groups of HS modifying enzymes was significantly downregulated by IFNγ+LPS, with significant changes in expression of genes that control HS chain elongation (*Ext2*, *Extl1*, *Extl2*), N-sulfation (*Ndst1*), 2-O-sulfation (*Hs2st1*), 3-O-sulfation (*Hs3st1*, *Hs3st3b1*) and 6-O-sulfation (*Hs6st1*, *Sulf2*) (Fig. 1A).

Polarisation with IL4+IL13 had limited effects, with altered expression of only three core proteins (increased *Gpc1*, *Sdc1*, *Sdc4* compared to unpolarised BMDM, Fig. 1A-C).

The most strongly regulated gene was *Sulf2*, a sulfatase that removes 6-O sulfate groups from N-acetylglucosamine residues of HS. Expression of this gene was markedly down-regulated in response to IFNγ+LPS treatment (25-fold compared to unpolarised BMDM, Fig. 1A).

### *Sulf2*-deficiency altered macrophage polarisation *in vitro*

We observed perinatal lethality in *Sulf2*^-/-^ mice, as has been reported by others [28, 29], so we utilised *Sulf2*^+/-^ BMDM to investigate the impact of partial *Sulf2* deficiency in macrophages. Given that unpolarised BMDMs expressed high levels of *Sulf2*, we investigated the effect of *Sulf2*-deficiency on polarisation of these cells (Fig. 2A). IL4+IL13-treated *Sulf2^+/-^* BMDMs expressed significantly lower levels of the anti-inflammatory markers *Arg1*, *Cd206* and *Fizz1*, indicating that *Sulf2*-deficiency reduced polarisation to an anti-inflammatory phenotype. IFNγ+LPS-treated *Sulf2^+/-^*BMDMs expressed significantly higher levels of the prototypic pro-inflammatory marker *Tnf* and reduced levels of *Socs3*, a suppressor of cytokine signaling (Fig. 2A), indicating a potentially more inflammatory phenotype, although these cells also expressed lower levels of *Nos2*.

**Fig. 2.**
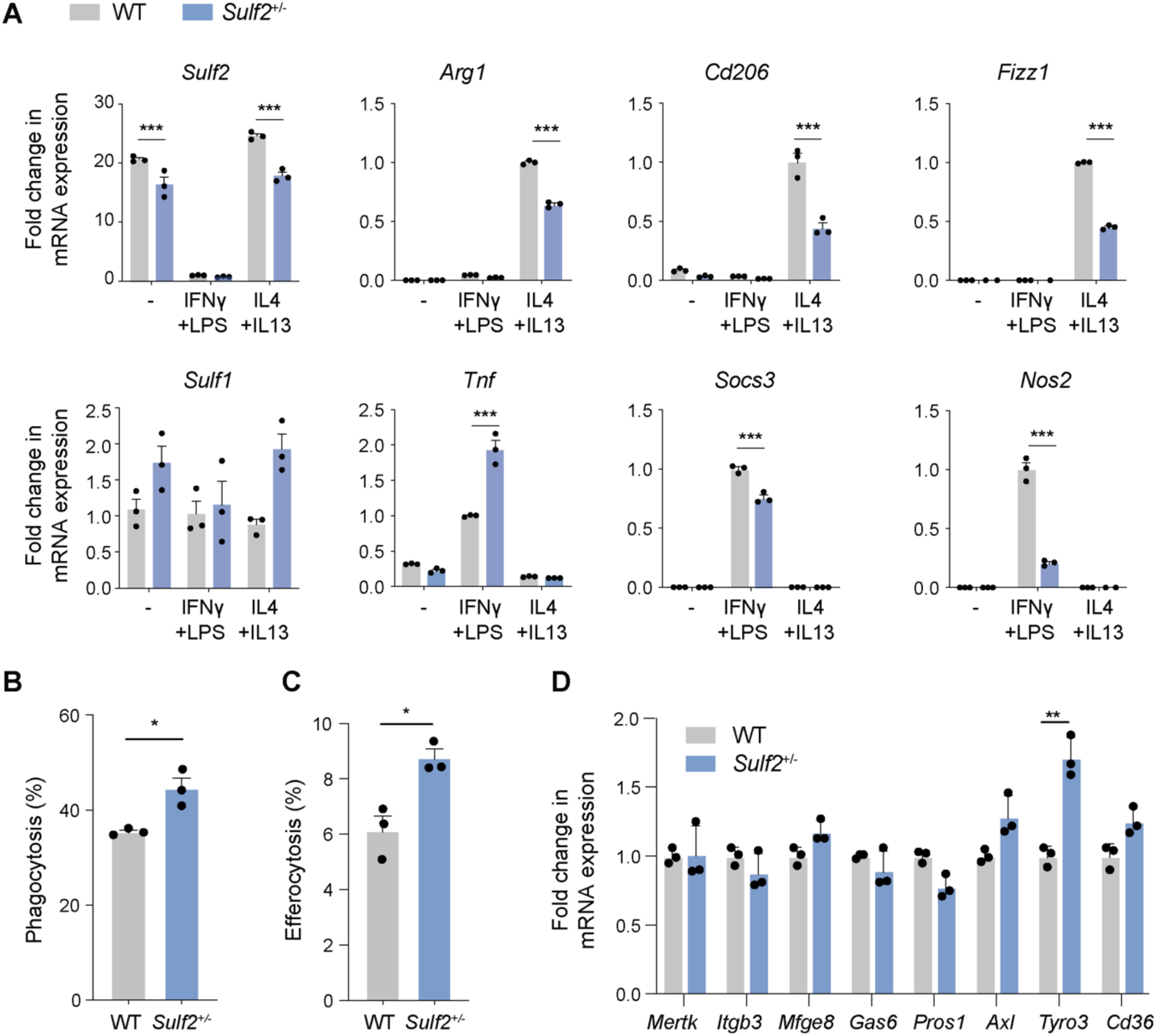
Macrophage polarisation and phagocytosis was modulated by *Sulf2*. BMDM from WT and *Sulf2^+/-^* mice were stimulated with IFNγ+LPS or IL4+IL13 (18 h). (**A**). Fold change in gene expression was quantified relative to *Rplp0* (n=3, mean ± SEM, analysed by two-way ANOVA with Bonferroni’s correction for multiple comparisons. ***, P<0.001). (**B-C**). Cells were incubated with fluorescent polystyrene beads at 50 MOI (B) or apoptotic cells at 1 MOI, and the percentage of percentage of fluorescence positive cells determined by flow cytometry (n=3, mean ± SEM, analysed by two-way Student’s *t*-tests. *, P<0.05). (**D**). Fold change in gene expression was quantified relative to *Rplp0* and to WT cells (n=mean ± SEM, analysed by multiple unpaired *t*-tests, followed by multiplicity correction using the two-stage step-up method of Benjamini, Krieger and Yekutieli, **, q<0.01).

Potential mechanisms leading to these changes in polarisation were investigated. There was no evidence for a change in IL6 release in response to IFNγ, LPS, IFNg+LPS, poly(I:C), FSL-1 or Pam3CSK4 (Fig. S1A), indicating TLR signaling was not impacted by *Sulf2*-deficiency. Similarly, there was no significant difference in pyroptotic cell death and IL1β release in response to LPS+nigerycin was unaffected (Fig. S1B), indicating unaltered inflammasome activation in *Sulf2*-deficient BMDM. HS has been shown to have effects on autophagy [30], but levels of LC3-II and autophagic flux were also unaltered in *Sulf2*-deficient BMDM (Fig. S1C).

### *Sulf2*-deficiency increased phagocytosis *in vitro*

To further assess the phenotype of *Sulf2*-deficient BMDM, we profiled their phagocytic and antigen-presenting abilities.

Mean phagocytosis of polystyrene beads was increased by 25% (Fig. 2B, P<0.05) and phagocytosis of apoptotic cells was increased by 43% (Fig. 2C, P<0.05) in *Sulf2*-deficient cells. This was accompanied by significantly increased expression of the efferocytosis receptor *Tyro3*, but not *Mertk*, *Axl* or *Cd36* (Fig. 2D). Expression of integrin αvβ3 (*Itgb3*) and the efferocytosis bridging molecules *Gas6* and *Pros1* was similarly unaffected.

*In vitro* uptake of OVA antigen and cell surface expression of costimulatory molecules (MHC-I, MHC-II, CD80, CD86) was not significantly altered in *Sulf2*-deficient BMDMs and DCs (Fig. S2). Similarly, there was no evidence that *Sulf2*-deficiency had an effect on *in vitro* presentation of OVA to CD4+ T cells by DCs, with unaltered CD4+ T cell proliferation and expression of CD25 (Fig. S3).

### Myeloid *Sulf2* deficiency increased joint damage and inflammation *in vivo*

*Sulf2* is highly expressed by myeloid cells (Fig. S4), but it is also expressed by multiple other cell types [31]. To study the function of myeloid *Sulf2*, we thus generated chimeric mice by irradiating WT animals and reconstituting their myeloid population with BMDM isolated from either WT or *Sulf2*^+/-^ donors. The effect of *Sulf2*-deficiency on myeloid function was then assessed using the antigen-induced arthritis (AIA) model, in which immunisation and boosting of mice with mBSA leads to myeloid and lymphoid cell-dependent joint inflammation and damage in mice.

WT and *Sulf2*^+/-^ chimeric mice developed similar acute inflammation in the first 4 days after intra-articular injection with mBSA, with no significant difference in initial paw swelling (Fig. 3A) or histological joint damage (Fig. S5). However, the resolution of inflammation was significantly impaired in *Sulf2*^+/-^ chimeric mice, with paw swelling significantly elevated at days 5-7. This was accompanied by increased joint damage in the *Sulf2*^+/-^ chimeric mice, with a significantly increased total histological score, synovial and subsynovial thickening and bone marrow density at day 7 (Fig. 3B-C). Paradoxically, *Sulf2*^+/-^ chimeric animals exhibited less pain at day 7 (Fig. 3C).

**Fig. 3.**
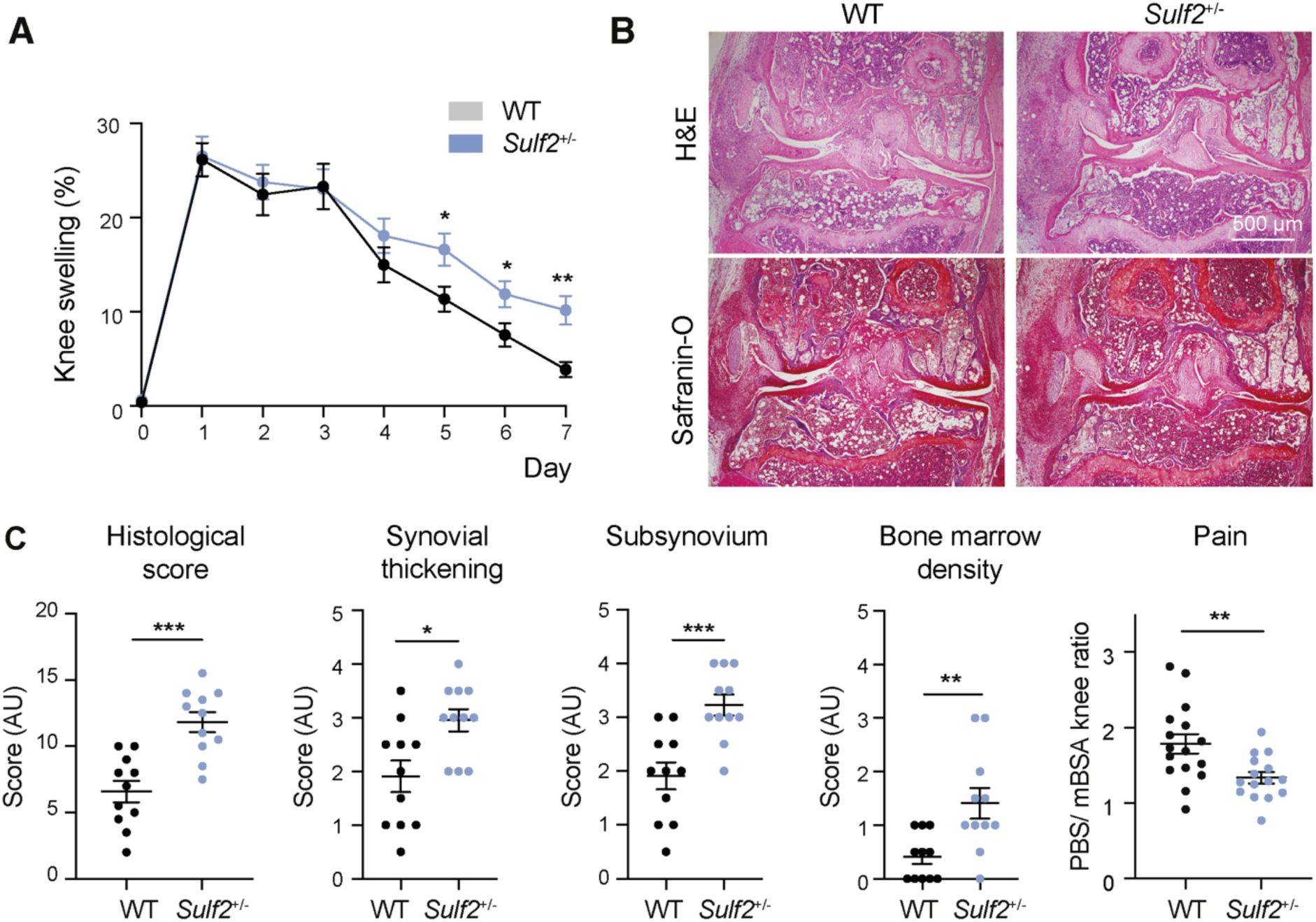
Swelling and AIA histology scores were increased at day 7 in *Sulf2*-deficient bone marrow chimera mice. WT and *Sulf2+/-* bone marrow chimeric mice (n=11-15) were immunised with mBSA in CFA (100 μg) and arthritis was induced 3 weeks later by intra-articular tibiofemoral injection of mBSA (100 μg, right knee). Mice were sacrificed 7 days later and disease severity in the knees was visualised histologically following safranin-O and H&E staining. (**A**). Knee swelling (n=15) was measured daily for 7 days after intra-articular injection using a calliper (mean ± SEM, analysed by a multiple unpaired *t*-tests, followed by multiplicity correction using the two-stage step-up method of Benjamini, Krieger and Yekutieli, **, q<0.01. *, q<0.05). (**B**). Representative histology at day 7. (**C**). Histology of the joint (4x magnification, n=11) was examined to calculate the total histological score, synovium and subsynovium thickness, and bone marrow density. Pain was assessed by measuring weight bearing using a Bioseb weight-bearing chamber, with data presented as the ratio of time spent on the PBS-injected limb relative to the mBSA-injected limb (mean ± SEM, analysed by a two-tailed Mann-Whitney u test. *, P<0.05; **, P<0.01; ***, P<0.001).

### Myeloid *Sulf2* deficiency increased Th17/Treg ratio in knees

The increased joint swelling in *Sulf2*^+/-^ chimeric mice was not explained by increased immune cell infiltration, as flow cytometry showed there was no significant difference in the abundance or frequency of myeloid or adaptive immune cell types in the knees of WT and *Sulf2*^+/-^ chimeric mice 2 or 7 days after initiation of AIA (Fig. S6).

While the overall number of CD4+ T cells was not altered (Fig. S6), closer examination of this compartment revealed a significant increase in Th17 abundance in the knees of *Sulf2*^+/-^ chimeric mice on day 7 of AIA (Fig. 4A-B), leading to a 2-fold increase in the Th17/Treg ratio (Fig. 4C-D). This shift in the Th17/Treg ratio was not observed in the inguinal lymph nodes (Fig. S7), indicating it was a local rather than a systemic effect.

**Fig. 4.**
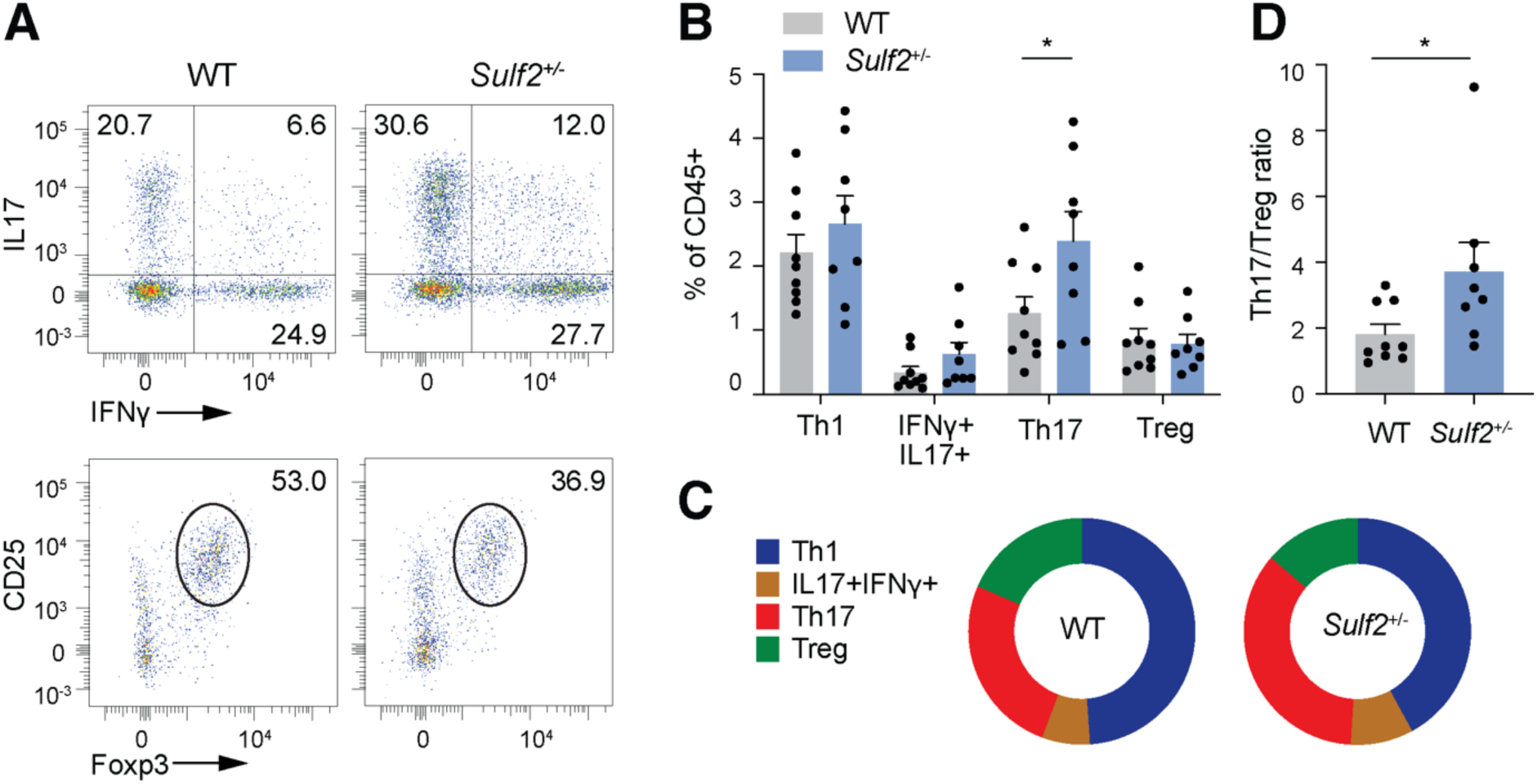
The Th17/Treg ratio was increased in the knee joints of *Sulf2*-deficient mice at day 7 of AIA. Single cell suspensions were isolated from knees of WT and *Sulf2^+/-^* bone marrow chimeric mice (n=9) 7 days after initiation of arthritis, and stimulated for 4 h with PMA (20 ng/ml) and ionomycin (1 mg/ml) in the presence of protein transport inhibitors. CD3+, CD4+ and CD8+ T-cell subsets were analysed by flow cytometry. (**A**). Representative dot plots for Th17 (IFNγ+IL17+) and Treg (CD25+Foxp3+) subsets in AIA knees. (**B**). The abundance of T cell subsets (as a percentage of CD45+ cells) was calculated (mean ± SEM, analysed by a two-tailed Student’s *t*-test. *, P<0.05). (**C**). Graphical representation of data in (**B**). (**D**). The Th17/Treg ratio was calculated from data in (**B**) (mean ± SEM, analysed by a two-tailed Student’s *t*-test. *, P<0.05).

### Type I interferon signaling was elevated in *Sulf2*-deficient macrophages

To gain insight into the mechanism underlying increased Th17 generation in *Sulf2*^+/-^ chimeric mice, we conducted bulk RNA sequencing on cells isolated from knees of WT and *Sulf2*^+/-^ chimeric mice 7 days after initiation of AIA (Fig. 5A). 199 genes were found to be significantly differentially expressed between WT and *Sulf2*^+/-^ chimeric mice (76 up-regulated and 123 down-regulated; Padj<0.05 by DESeq2, Table S1). Deconvolution of the data with MuSIC [32] estimated that the vast majority of sequences (>85%) originated from macrophages (Fig. S8). KEGG pathway enrichment analysis of the significantly differentially expressed genes was indicative of altered type I interferon signaling, with significantly enriched KEGG identifiers including “Herpes simplex virus infection 1” (P=1.4×10^-4^, gene ratio=0.16), “Hepatitis B” (P = 3.2×10^-3^, gene ratio=0.07) and “Human T-cell leukemia virus 1 infection” (P = 3.6×10^-3^, gene ratio=0.07)(Fig. 5B). Subsequent comparison with the Interferome database [33] indicated that 111 of the 199 differentially expressed genes (56 %) are known to be type I interferon-responsive genes (marked in red on Fig. 5A). Analysis of the promoter region of differentially regulated genes indicated that 42% contain recognition sites for the type I interferon-related transcription factor ELF4 [34] (p = 1×10^-2^) and 11% contain recognition sites for IRF3 [35] (p = 1×10^-2^), both transcription factors known to bind to interferon-stimulated response elements.

**Fig. 5.**
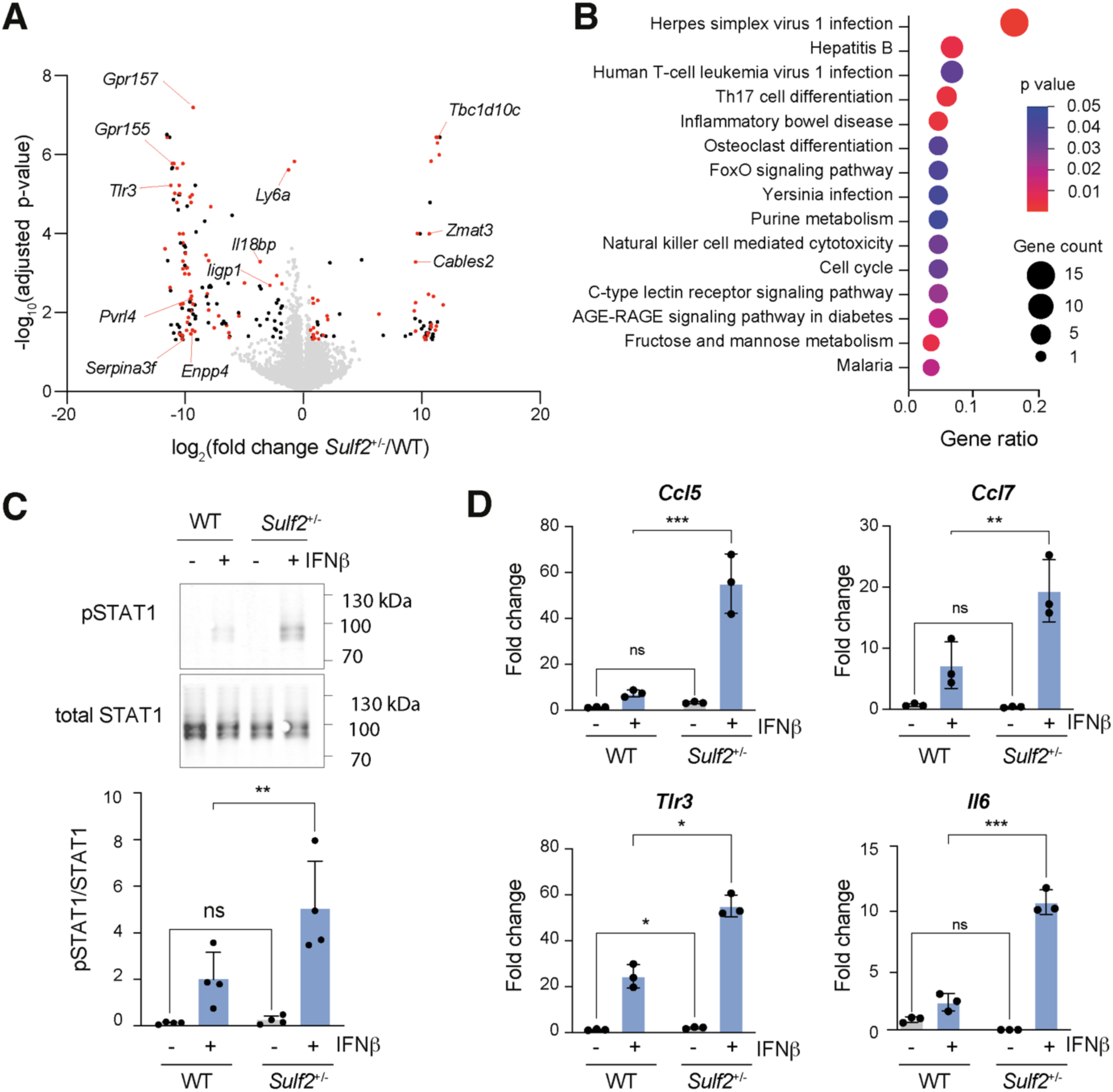
Type I interferon signaling was increased in in *Sulf2*-deficient macrophages. (**A-B**). Single cell suspensions were isolated from knees of WT and *Sulf2^+/-^*bone marrow chimeric mice (n=3) 7 days after initiation of arthritis and bulk RNA sequencing performed. (**A**). The volcano plot shows differentially expressed genes (fold change in *Sulf2*^+/-^ compared to WT) identified by DeSeq2, with significantly altered type I interferon-responsive genes[33] labelled in red, other significantly altered genes in black, and non-significantly altered genes in grey. (**B**). The top 15 KEGG identifiers enriched in differentially expressed genes are shown, indicating their P-value and gene ratio (fraction of the differentially-expressed genes in each KEGG identifier). (**C**). BMDM from WT and *Sulf2^+/-^* mice were stimulated with IFNβ (50 ng/ml, 30 min) and phosphorylation of STAT1 quantified by immunoblotting (n=4 biological replicates per group, mean ± SD, analysed by a 2-way ANOVA with Sidak’s correction for multiple comparisons. **, P<0.01**). (**D**). WT and *Sulf2^+/-^* BMDM (n=3 biological replicates per group) were stimulated with IFNβ (50 ng/ml, 4 h) and expression of *Ccl5*, *Ccl7*, *Tlr3* and *Il6* quantified by RT-qPCR relative to *Gapdh* (n=3 biological replicates per group, mean ± SD, analysed by a 2-way ANOVA with Sidak’s correction for multiple comparisons. *, P<0.05; **, P<0.01; ***, P<0.001).

To validate these findings, WT and *Sulf2*^+/-^ BMDM were stimulated *in vitro* with IFNβ and phosphorylation of STAT1 quantified. This showed a significant increase in pSTAT1 in *Sulf2*^+/-^ BMDM (Fig. 5C). Similarly, *Sulf2*^+/-^ BMDM expressed significantly higher levels of the interferon-stimulated genes *Ccl5*, *Ccl7*, *Tlr3* and *Il6* in response to IFNβ stimulation (Fig. 5D).

## Discussion

### HS biosynthesis is significantly regulated during macrophage polarisation

Both the immune system and HS proteoglycans are complex information networks, in which dynamic interactions between multiple nodes determine biological outcomes. Here, we comprehensively profiled changes in expression of HS biosynthetic machinery during macrophage polarisation, and found that almost two-thirds of the expressed genes were significantly regulated in response to IFNγ+LPS and/or IL4+IL13, with significant changes in expression of at least one gene from each family of HS modifying enzymes. The most highly regulated gene was the extracellular HS 6-O-sulfatase *Sulf2*, which was downregulated 25-fold by IFNγ+LPS. This extends previous work by Martinez *et al*. [23], who demonstrated significant changes in expression of HS sulfotransferases during macrophage polarisation, but did not examine HS polymerases (*Ext1*, *Ext2*, *Extl1*, *Extl2*, *Extl3*), epimerase (*Glce*), degrading enzymes (*Hpse*, *Hpse2*) or sulfatases (*Sulf1*, *Sulf2*).

The strong regulation of *Sulf2* expression is striking because SULF1 and SULF2 are the only enzymes that are able to modify HS sulfation in the extracellular environment [11]. N-, 2-O-, 3-O- and 6-O-sulfation are all added to HS during its synthesis in the Golgi apparatus, but once HS is secreted into the extracellular environment, its structure can only be modified by heparanase (encoded by *Hspe*), which cleaves HS into smaller fragments, or by the sulfatases (encoded by *Sulf1* and *Sulf2*), which remove sulfate groups specifically from the 6-carbon position of N-acetylglucosamine residues [31]. SULFs thus carry out the only ‘editing’ of HS that occurs in the extracellular environment, giving these enzymes the unique ability to rapidly fine-tune HS affinity for ligands and so dynamically modulate downstream signaling pathways [36].

### *Sulf2*-deficient macrophages have an elevated inflammatory phenotype

The marked reduction in *Sulf2* expression in response to treatment with IFNγ+LPS suggests that SULF2 may have an anti-inflammatory or regulatory role in macrophages. We found that *Sulf2*^+/-^ macrophages exhibited a generally more inflammatory phenotype *in vitro*, with higher expression of *Tnf*, and lower expression of *Socs3*, *Arg1*, *Cd206* and *Fizz* in response to polarising cytokines. This is in agreement with Zhang *et al*. [37], who showed that loss of SULF2 expression in bladder cancer cells promoted polarisation of co-cultured THP-1 cells towards an inflammatory phenotype. Further *in vitro* investigations showed that no evidence that macrophage *Sulf2*-deficiency altered TLR signaling, inflammasome activation, autophagy or antigen presentation, so we sought to investigate the role of myeloid *Sulf2* on inflammation *in vivo*.

*Sulf2* is widely expressed [31], so we adopted a bone marrow transfer approach, to generate chimeric mice with heterozygous deficiency of *Sulf2* in the myeloid lineage. We observed perinatal lethality of *Sulf2*^-/-^ animals, as has been reported for some other *Sulf2*^-/-^ lines [28, 29], but not others [38, 39]. Such strain-dependent effects have also been reported for mice lacking other HS biosynthetic enzymes, such as *Hs3st1* [40] and *Hs6st1* [41], suggesting HS biosynthesis is strongly influenced by modifier genes.

*Sulf2*^+/-^ bone marrow chimeric mice showed significantly increased joint swelling and histological damage in the resolution phase of the macrophage-[42, 43]- and CD4^+^ T cell-dependent [44] antigen-induced arthritis model. This was accompanied by an increased abundance of Th17 cells in joints of *Sulf2*^+/-^ chimeras, consistent with the known ability of this subset to exacerbate inflammation in this [45] and other [46] models of arthritis.

### Increased type I interferon signaling in *Sulf2*-deficient macrophages

To understand how *Sulf2*-deficiency in macrophages could promote Th17 differentiation, we conducted bulk RNA sequencing of cells isolated from AIA joints during the resolution phase of joint inflammation. This showed that differentially expressed genes were related to viral infection responses, suggesting altered type I interferon signaling, with 56% of the differentially expressed genes being known type I interferon-responsive genes. *In vitro* analyses confirmed elevated responses to IFNβ in *Sulf2*^+/-^ BMDM, with increased STAT1 phosphorylation and increased induction of *Ccl5*, *Ccl7*, *Tlr3* and *Il6* expression. Increased IL-6 expression would lead to the observed increase in Th17 abundance and Th17/Th1 ratio [47]. This proposed mechanism is consistent with a previous study in lung carcinoma cells, where siRNA or epigenetic silencing of SULF2 was found to activate expression of interferon-inducible genes [48].

Considering the molecular mechanism by which SULF2 could reduce type I interferon signaling, Gordts *et a*l. [26] previously showed that IFNβ signaling in macrophages is modulated by cell surface HS. IFNβ binds strongly to HS (*K*i of 1.4 nM) [26, 49], while IFNα4, interferon-alpha/beta receptor alpha chain (IFNAR1) and interferon-alpha/beta receptor beta chain (IFNAR2) do not [26], and as with all HS ligands, IFNβ/HS affinity is likely to be modulated by the overall level and pattern of HS sulfation. N-sulfation of HS was shown to reduce IFNβ signaling and to be protective in models of atherosclerosis and obesity [26]. Our data here show that 6-O-sulfation of HS has an opposite effect, promoting IFNβ signaling and leading to increased *Il6* expression, which promotes the generation of Th17 cells and exacerbates inflammatory arthritis.

We propose a model in which N-sulfation of HS promotes IFNβ binding to cell surface HS proteoglycans in preference to its cognate receptors, while 6-O-sulfation generates a different structural motif that promotes formation and/or stabilisation of IFNβ/receptor complexes (Figure 6). Homeostatic expression of SULF2 would keep 6-O-sulfation of cell surface HS at low levels, leading to accumulation of IFNβ on N-sulfated HS proteoglycans and keeping IFNβ signaling in check. In inflammatory environments (or *Sulf2*-deficiency, which mimics and/or exacerbates this), reduced SULF2 would lead to increased HS 6-O-sulfation and elevated IFNβ signaling. This predicts that IFNβ signaling would be further elevated in macrophages deficient for both *Sulf2* and *Ndst1*.

**Fig. 6.**
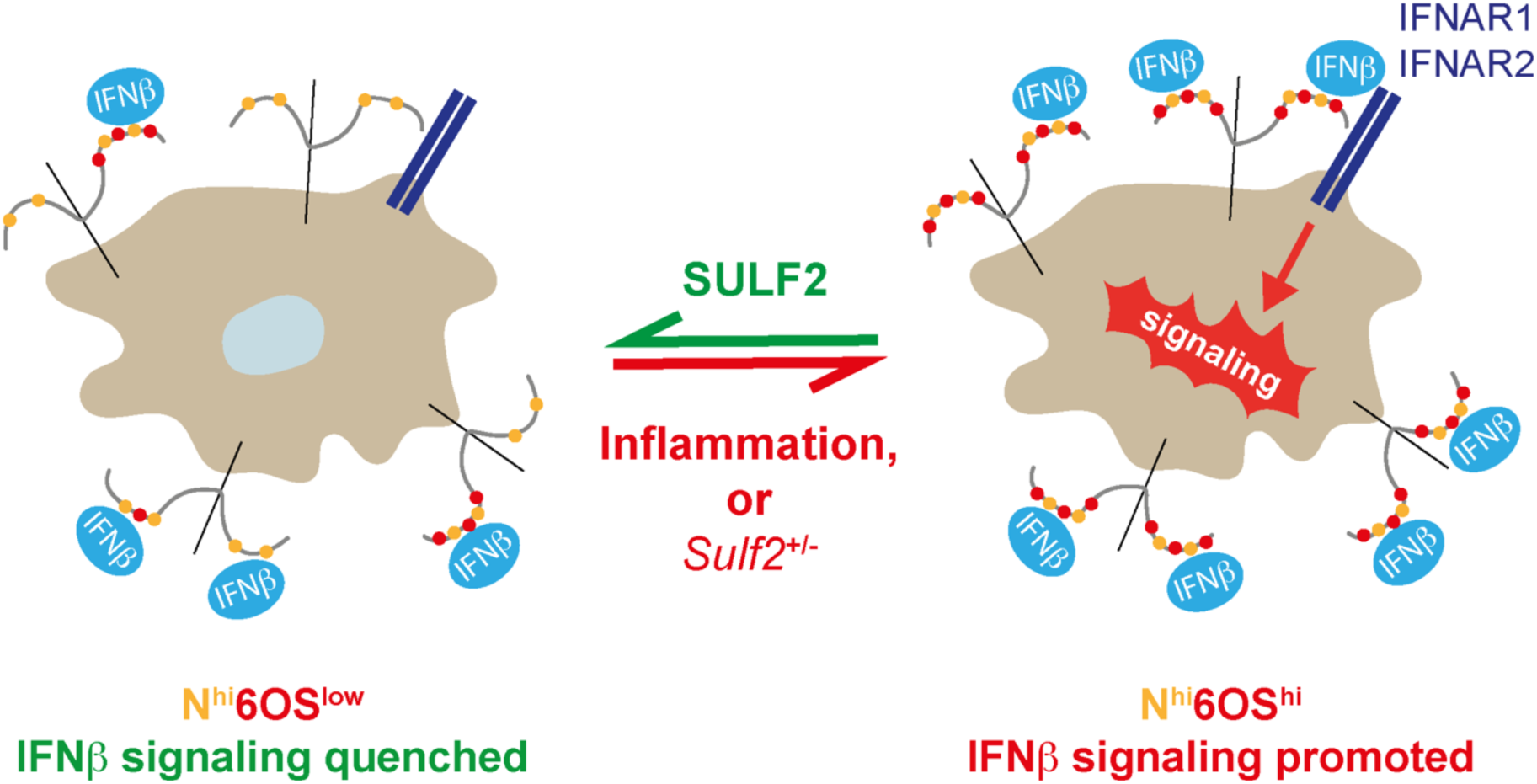
Proposed model of HS regulation of IFNΒ signaling. N-sulfation of HS promotes IFNβ binding to cell surface HS proteoglycans in preference to engagement with IFNAR receptors [26], while 6-O-sulfation promotes formation and/or stabilisation of IFNβ/receptor complexes. High expression of NDST1/2 [26] and HS6ST1 would enable macrophages to synthesise highly N- and 6-O-sulfated HS, but homeostatic expression of SULF2 would generate an N^hi^6OS^low^ motif on the cell surface, favouring IFNβ sequestration. In inflammatory environments (or *Sulf2*-deficiency, which mimics and/or exacerbates this), reduced SULF2 activity would allow an N^hi^6OS^hi^ HS motif to accumulate, promoting IFNβ signaling. SULF2 thus serves to limit inflammatory IFNβ signaling in macrophages.

*Sulf2* is thus likely to also be protective in other autoimmune conditions where IFNβ signaling promotes Th17-driven inflammation e.g. psoriasis, systemic lupus erythematosus, Sjögren’s syndrome, and neuromyelitis optica [50]. However, IFNβ signaling can also suppress auto-immune responses, such as in relapsing remitting multiple sclerosis, where IFNβ is used therapeutically to slow disease progression[50], and *Sulf2* may have deleterious effects in such contexts. Indeed, Saraswat *et al*. [51] showed that *Sulf2* expression is elevated in multiple sclerosis lesions, and propose that it contributes to an inhibitory microenvironment that limits remyelination. HS sulfation in numerous tissues and cells types is known to change with ageing [13–15], inflammation [52], infection [53] and metabolic state [54, 16], suggesting a novel mechanism by which co-morbidities could modulate physiological, pathological and therapeutic type I interferon signaling. Establishing what regulates *Sulf2* expression and HS structure more broadly is thus likely to be of importance for multiple auto-immune conditions. **Further considerations**. Looking beyond type I interferon signaling, *Sulf2*-deficiency is also likely to impact the biological activity of multiple other HS-binding proteins, with experimental identification of an affected pathway dependent on the model analysed and its molecular drivers. For example, *Sulf2*-deficient mice have worse outcomes in models of osteoarthritis [55], myocardial infarction [56], and bleomycin-induced lung injury [57], which has been attributed to impaired BMP [55] and VEGF [56] signaling and elevated FGF2 [55] and TGFβ [57] signaling.

Both cell-associated [58–60] and secreted [31, 56, 59] forms of SULF2 have been described, suggesting the enzyme can act in both an autocrine and paracrine manner. Recently, El Masri *et al*. [61] found that SULF2 localisation and activity can be modulated by post-translational modification, with attachment of a chondroitin or dermatan sulfate glycosaminoglycan chain to the enzyme’s HD domain increasing cell surface association and reducing 6-O-sulfatase activity. Degradation of the glycosaminoglycan chain by hyaluronidase reduced cell surface binding and increased SULF2 activity [61]. Conditions that increase hyaluronidase expression could thus switch SULF2 from autocrine to enhanced paracrine activity. Our *in vivo* data are likely to reflect autocrine activity of SULF2, because *Sulf2*-deficiency in myeloid cells was not compensated for by expression in other cells of the joint, such as synovial fibroblasts, which express abundant *Sulf2* in RA [62]. Intriguingly, Siegel *et al*. [62] found that SULF2 expressed by rheumatoid synovial fibroblasts promotes inflammation, by increasing transcriptomic and phenotypic responses of these cells to TNF. This suggests that synovial fibroblast and myeloid SULF2 have opposing effects on joint inflammation, with synovial SULF2 potentially contributing to initiation of inflammation, and myeloid SULF2 promoting resolution. Further investigation is required to define the kinetics of SULF2 expression and activity in these cell types.

## Author contributions

Conceptualization: MS, LT. Methodology: MS, AR, RC, AC, LT. Investigation: MS, AR, JO, RC, LT, EBC, MG, AC, LT. Resources: RW, DSB, NS, CM, LT. Discussion and advice: AR, JO, AC, RW, NS, CM, LT. Writing – original draft: MS, LT. Writing – review and editing: AR, RC, AC, RW, DSB, NS, LT. Funding acquisition: DSB, NS, CM, LT. Supervision: RW, DSB, NS, CM, LT. All authors read and approved the final manuscript.

## Data availability

The datasets used and analysed during the current study are available from the corresponding author on reasonable request.

## Declarations

### Ethical approval

All animal experiments were conducted in accordance with the United Kingdom Home Office Animals Scientific Procedures Act 1986 under relevant personal and project licenses.

### Consent to participate and consent to publish

Not applicable.

## Acknowledgements

None.

## Abbreviations

AIA: antigen-induced arthritis
APRIL: A Proliferation-Inducing Ligand
BMDM: bone marrow-derived macrophages
DCs: dendritic cells
FBS: fetal bovine serum
FSL-1: fibroblast-stimulating lipopeptide-1
FMO: fluorescence minus one
GM-CSF: granulocyte-macrophage colony-stimulating factor
HS: heparan sulfate
IFN: interferon
IFNAR1: interferon-alpha/beta receptor alpha chain
IFNAR2: interferon-alpha/beta receptor beta chain
KEGG: Kyoto Encyclopedia of Genes and Genomes
LC3: microtubule-associated protein light chain 3
LPS: lipopolysaccharide
mBSA: methylated bovine serum albumin
M-CSF: macrophage colony-stimulating factor
MFI: mean fluorescence intensity
MOI: multiplicity of infection
OVA: ovalbumin
Pam3CSK4: N-palmitoyl-S-[2,3-bis(palmitoyloxy)-(2RS)-propyl]-[R]-cysteinyl-[S]-seryl-[S]-lysyl-[S]-lysyl-[S]-lysyl-[S]-lysine
PCA: principal component analysis
PMA: phorbol 12-myristate 13-acetate
polyinosinic:polycytidylic acid: poly (I:C)
RPMI: Roswell Park Memorial Institute 1640 medium
SULF2: sulfatase 2
TaqMan Low-Density Array: TLDA
TLR: Toll-like receptor transforming growth factor beta (TGFβ)
WT: wild-type

---

**Supplementary Fig. S1.**
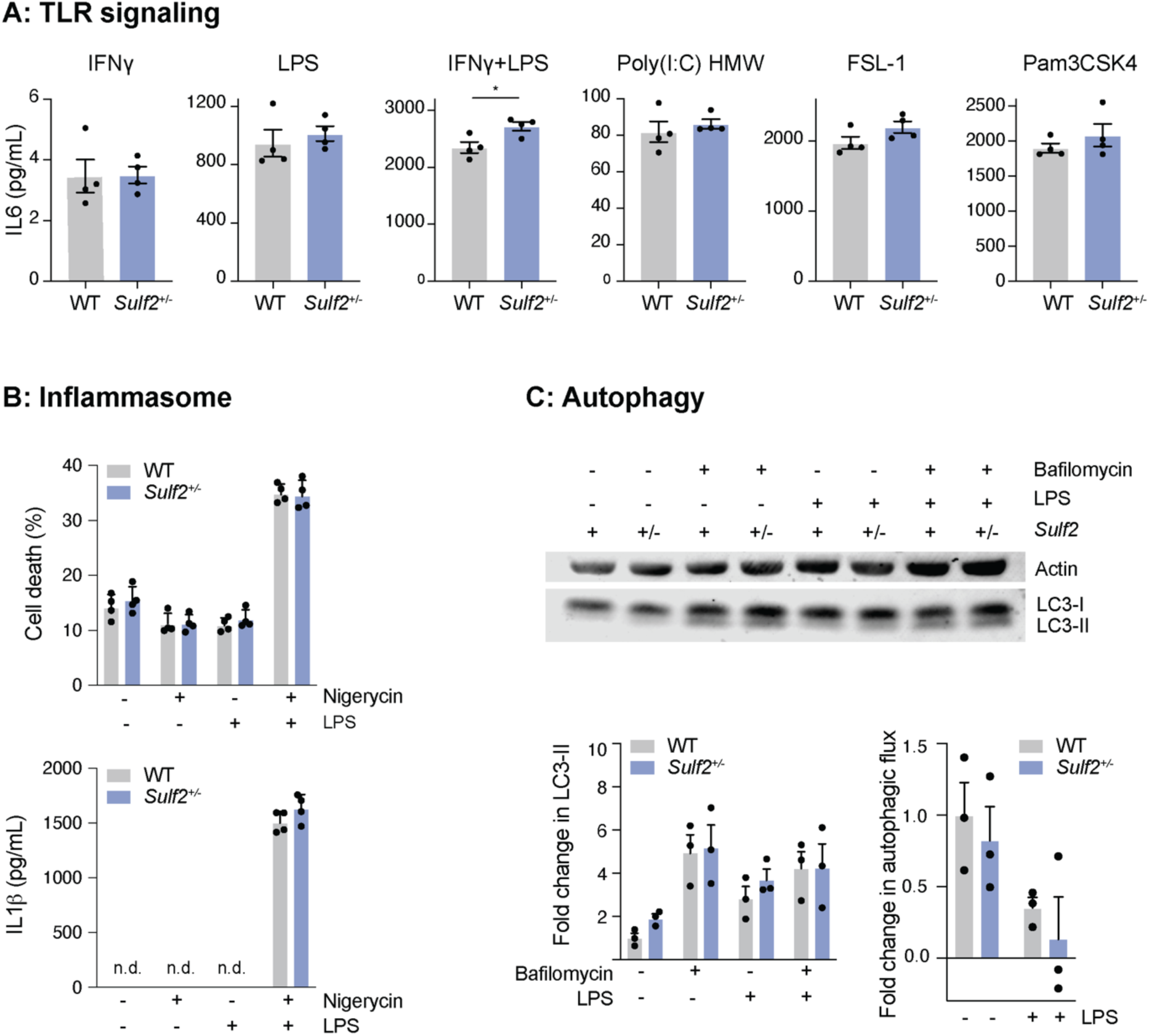
TLR signaling, inflammasome activation and autophagy were unchanged in *Sulf2*^+/-^ BMDM. (**A**). BMDM from WT and *Sulf2^+/-^* mice were stimulated for 6 h with IFNγ, LPS, IFNγ+LPS (all at 100 ng/ml), FSL-1 (100 ng/ml), poly(I:C) (10 ng/ml) or Pam3CSK4 (100 ng/ml) and IL6 secretion was measured by ELISA (n=4, mean ± SEM, analysed by a two-tailed Student’s T-test. *, P<0.05). (**B**). BMDM from WT and *Sulf2^+/-^* mice were stimulated with LPS (100 ng/ml, 6 h) with nigericin (5 μM) added for the last 2 h of culture. Cell death and IL1β secretion were quantified (n=4, mean ± SEM, analysed by two-way ANOVA with Bonferroni’s correction for multiple comparisons). (**C**). BMDM from WT and *Sulf2^+/-^* mice were stimulated with LPS (100 ng/ml, 18 h) and then treated with 100 nM bafilomycin (100 nM, 2 h) before LC3-I and LC3-II levels were quantified by immunoblotting. Fold change in LC3-II normalised to actin and the fold change in autophagic flux (LC3II in bafilomycin – untreated) normalised to actin were calculated (n=3, mean ± SEM, analysed by two-way ANOVA with Bonferroni’s correction for multiple comparisons).

**Supplementary Fig. S2.**
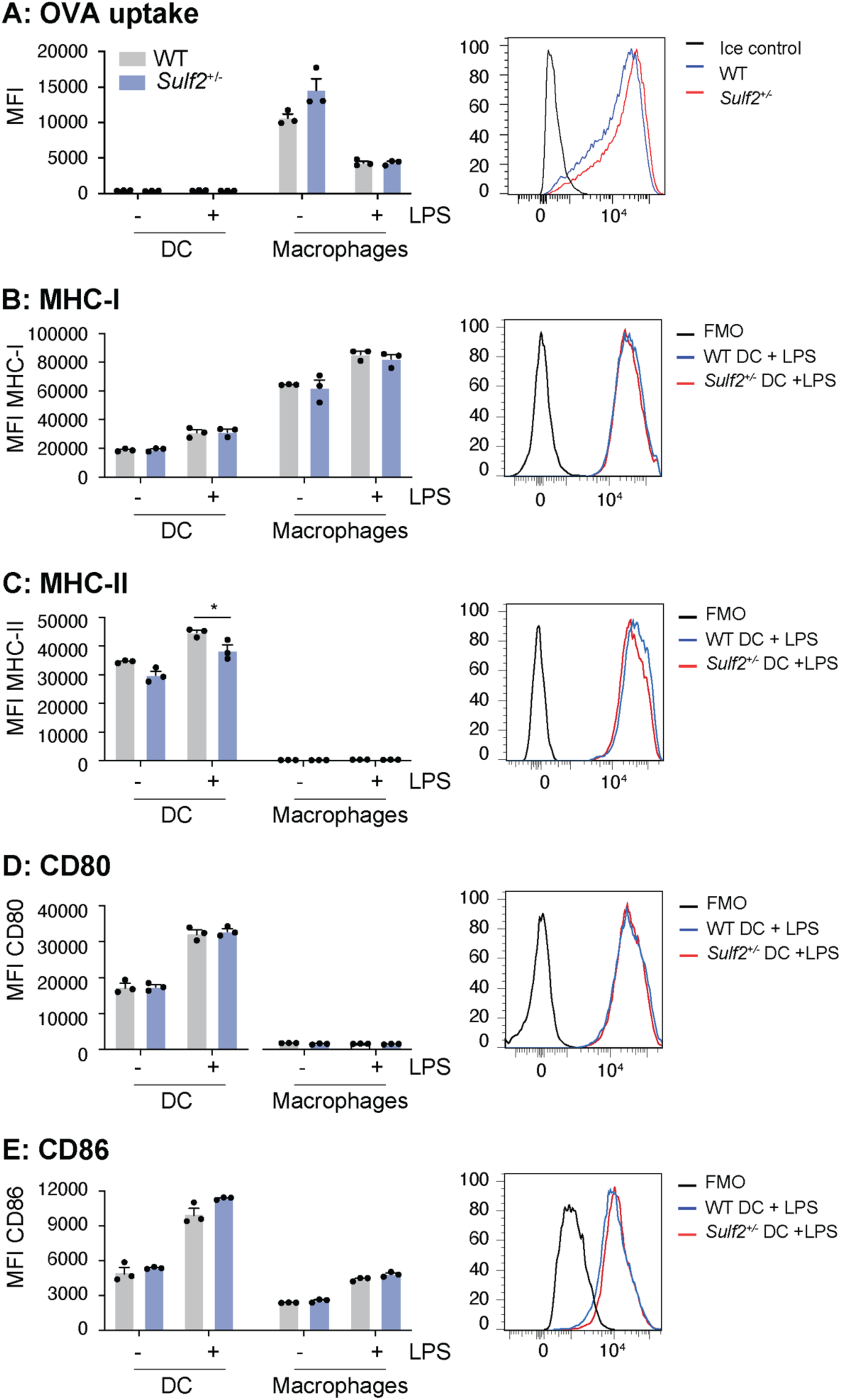
OVA uptake and expression of co-stimulatory molecules was unchanged in *Sulf2*^+/-^ macrophages. Bone marrow cells from WT and *Sulf2^+/-^* mice (n=3) were cultured with M-CSF (100 ng/ml, 7 d) to generate BMDM, or with GM-CSF (20 ng/ml, 7 d). (**A**). Cells were stimulated with LPS (100 ng/ml, 6 h) and incubated with Alexa647-conjugated OVA (0.1 mg/ml, 1 h, 37 °C) before analysis by flow cytometry. DCs in the GM-CSF-generated cell population were analysed by gating for CD11c+MHC-II^int-high^, followed by gating for CD11b^int^MHC-II^high^ (n=3, mean ± SEM, analysed by a two-way ANOVA with Bonferroni’s correction for multiple comparisons; *, P<0.05). (**B-E**). Cells were stimulated with LPS (100 ng/ml, 6 h) and cell surface expression of MHC-I (**B**), MHC-II (**C**), CD80 (**D**) and CD86 (**E**) measured by flow cytometry using the gating strategy described in (**A**) (n=3, mean ± SEM, analysed by a two-way ANOVA with Bonferroni’s correction for multiple comparisons; *, P<0.05).

**Supplementary Fig. S3.**
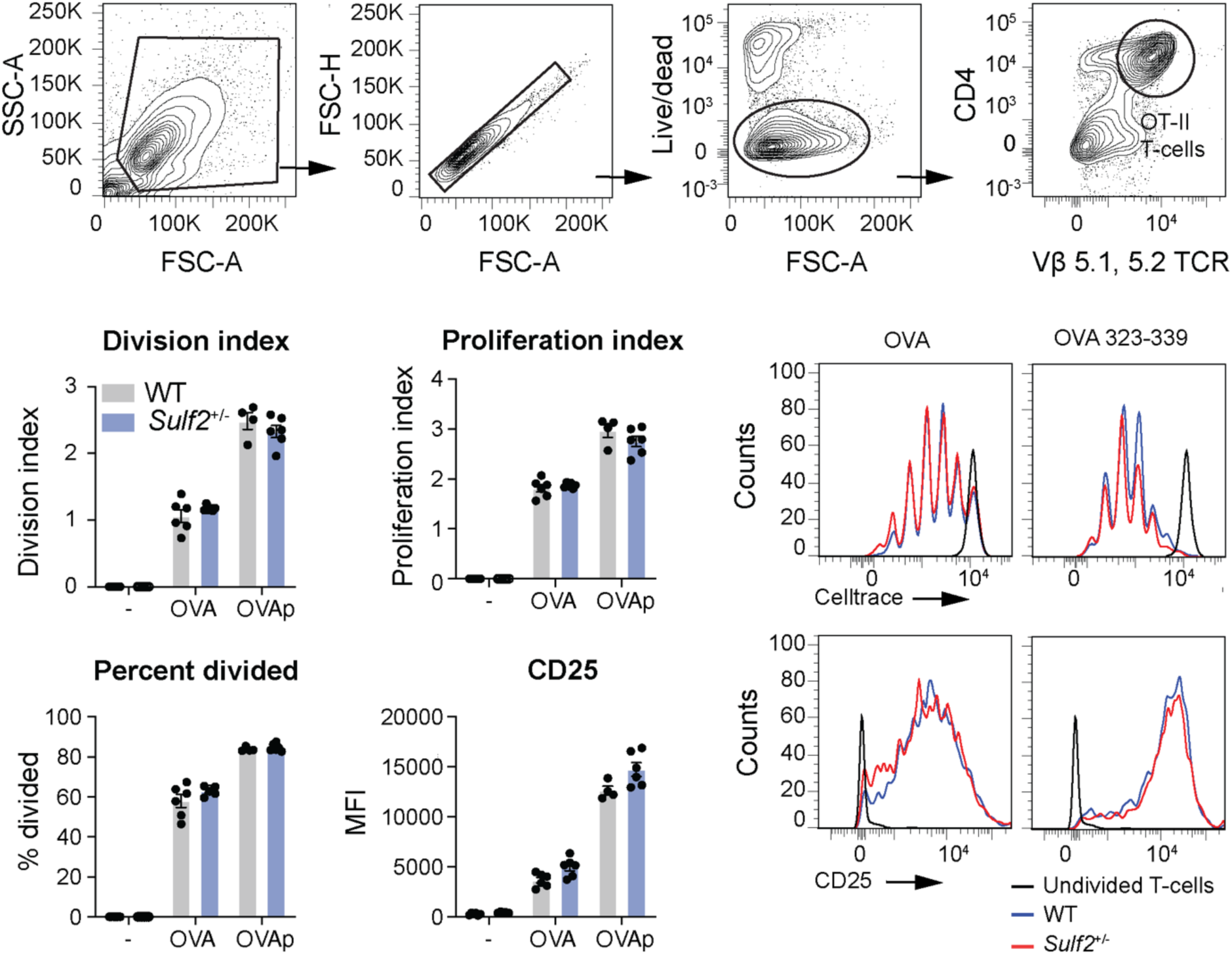
*Sulf2*-deficiency did not affect dendritic cell presentation of OVA antigen *in vitro*. CD4+ OT-II T-cells were labelled with CellTrace Violet and co-cultured (4 d) with LPS-activated WT or *Sulf2*^+/-^ DCs. OVA was added to antigen-presenting cells before LPS-activation, or OVA peptide 323-339 was added during co-culture with CD4+ T-cells. CD4+ T-cell proliferation and expression of CD25 were analysed by flow cytometry, using the gating strategy shown in top panel (n=3, analysed in duplicate, mean ± SEM, analysed by a two-way ANOVA with Bonferroni’s correction for multiple comparisons).

**Supplementary Fig. S4.**
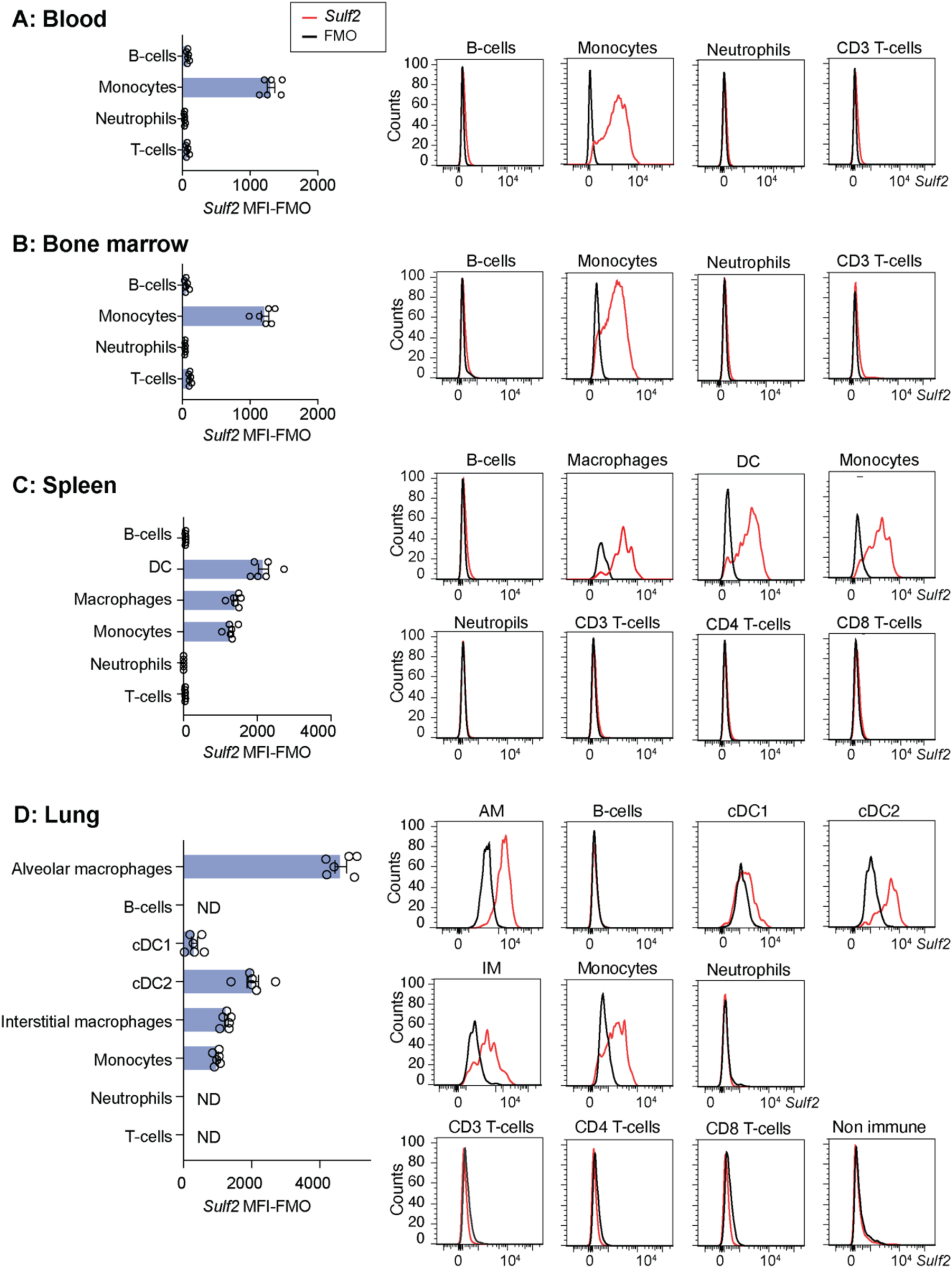
*Sulf2* was highly expressed by monocytes in murine blood and bone marrow, and by macrophages, monocytes and dendritic cells in the spleen and lung. *Sulf2* expression in immune populations was quantified by analysing single cell suspensions from blood, bone marrow, spleen and lung of C57BL/6 mice using the PrimeFlow RNA assay. Fluorescence was analysed by flow cytometry and expressed as the mean fluorescence intensity (MFI) minus the *Sulf2* fluorescence minus one (FMO). Representative histograms MFI-MFO (± SEM, n=6) are shown.

**Supplementary Fig. S5.**
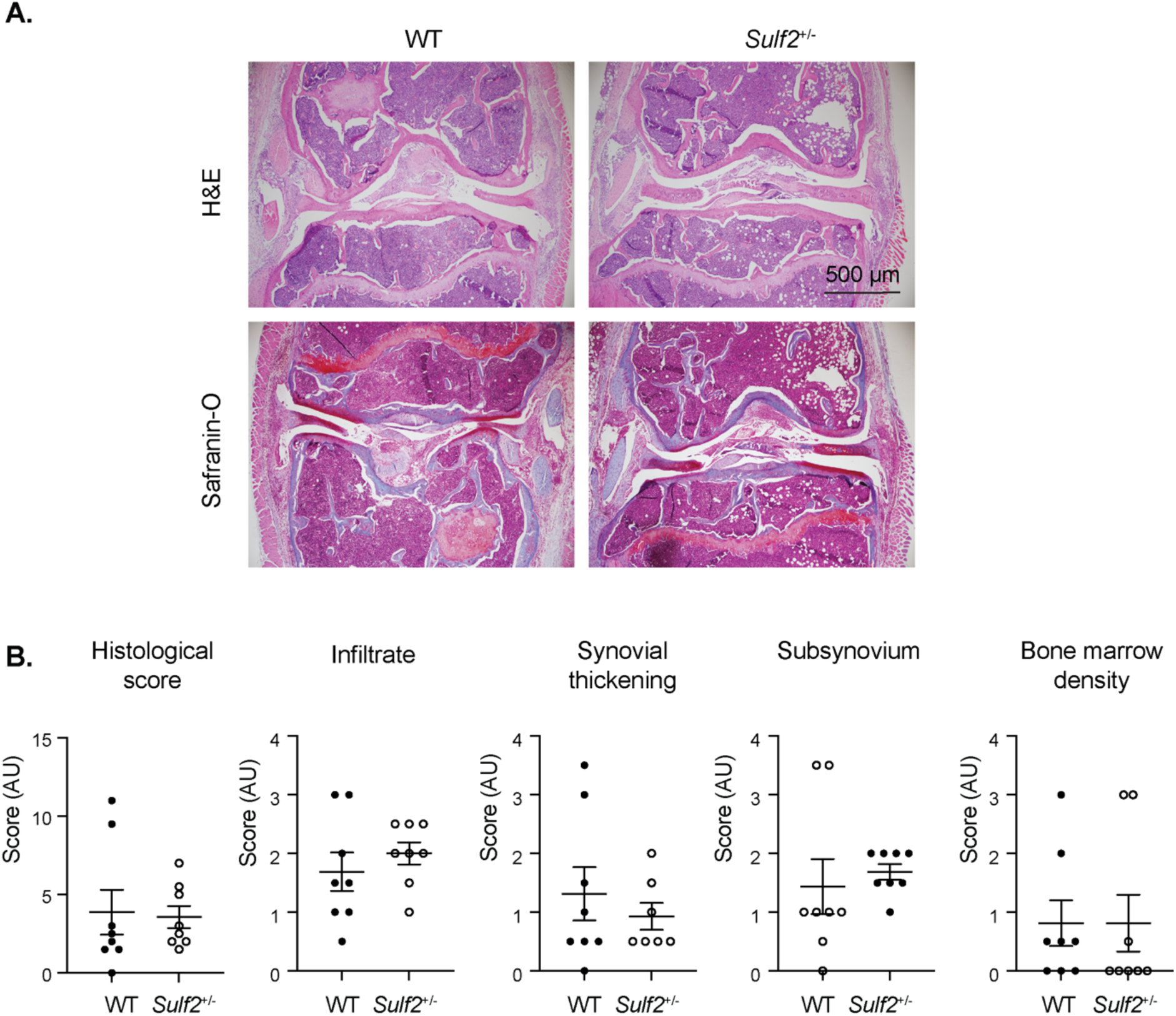
AIA histology scores were not affected at day 2 in *Sulf2*-deficient bone marrow-chimera mice. (**A**). WT and *Sulf2^+/-^* bone marrow chimeric mice (n=8) were immunised with mBSA (100 μg) in CFA, and arthritis induced 3 weeks later by intra-articular tibiofemoral injection of mBSA (100 μg, right knee). 2 days later, the mice were sacrificed and sections of the knees stained with safranin-O and H&E. (**B**). Histology of the joint (4x magnification) was examined to calculate the total histological score, synovium and subsynovium thickness, and bone marrow density (mean ± SEM, analysed by a two-tailed Mann-Whitney u test).

**Supplementary Fig. S6.**
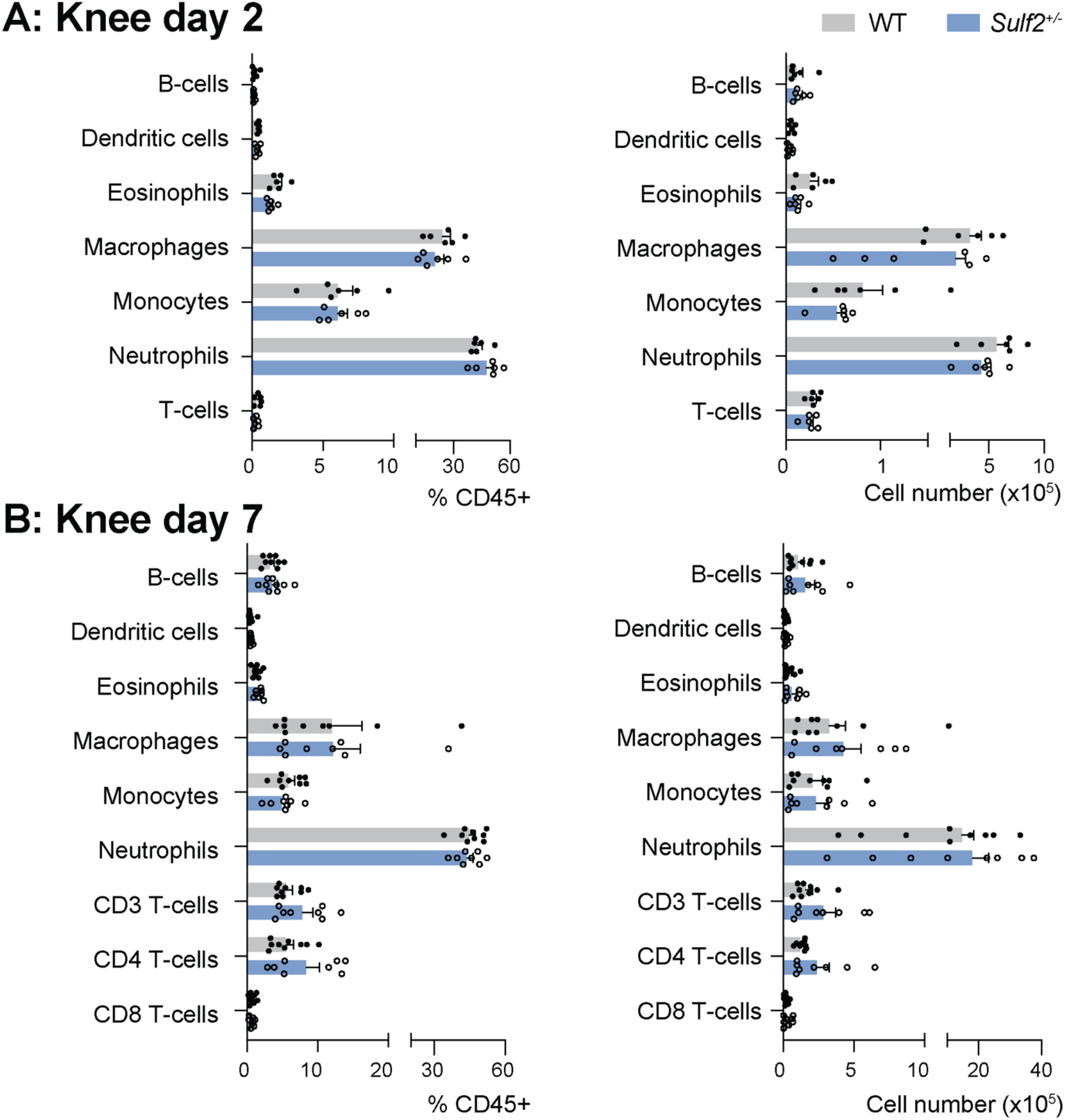
*Sulf2*-deficiency did not alter the abundance or frequency of immune cell types in the knee of AIA mice. WT and *Sulf2^+/-^* bone marrow chimeric mice were immunised with mBSA (100 μg) and arthritis was induced 3 weeks later by intra-articular tibiofemoral injection of 100 μg mBSA (right knee). 2 days (**A**, n=6) or 7 days (**B**, n=9) later, affected knees were digested and the prevalence of myeloid and adaptive immune cell populations was determined by flow cytometry (mean ± SEM, analysed by a multiple analysed by multiple unpaired *t*-tests, followed by multiplicity correction using the two-stage step-up method of Benjamini, Krieger and Yekutieli).

**Supplementary Fig. S7.**
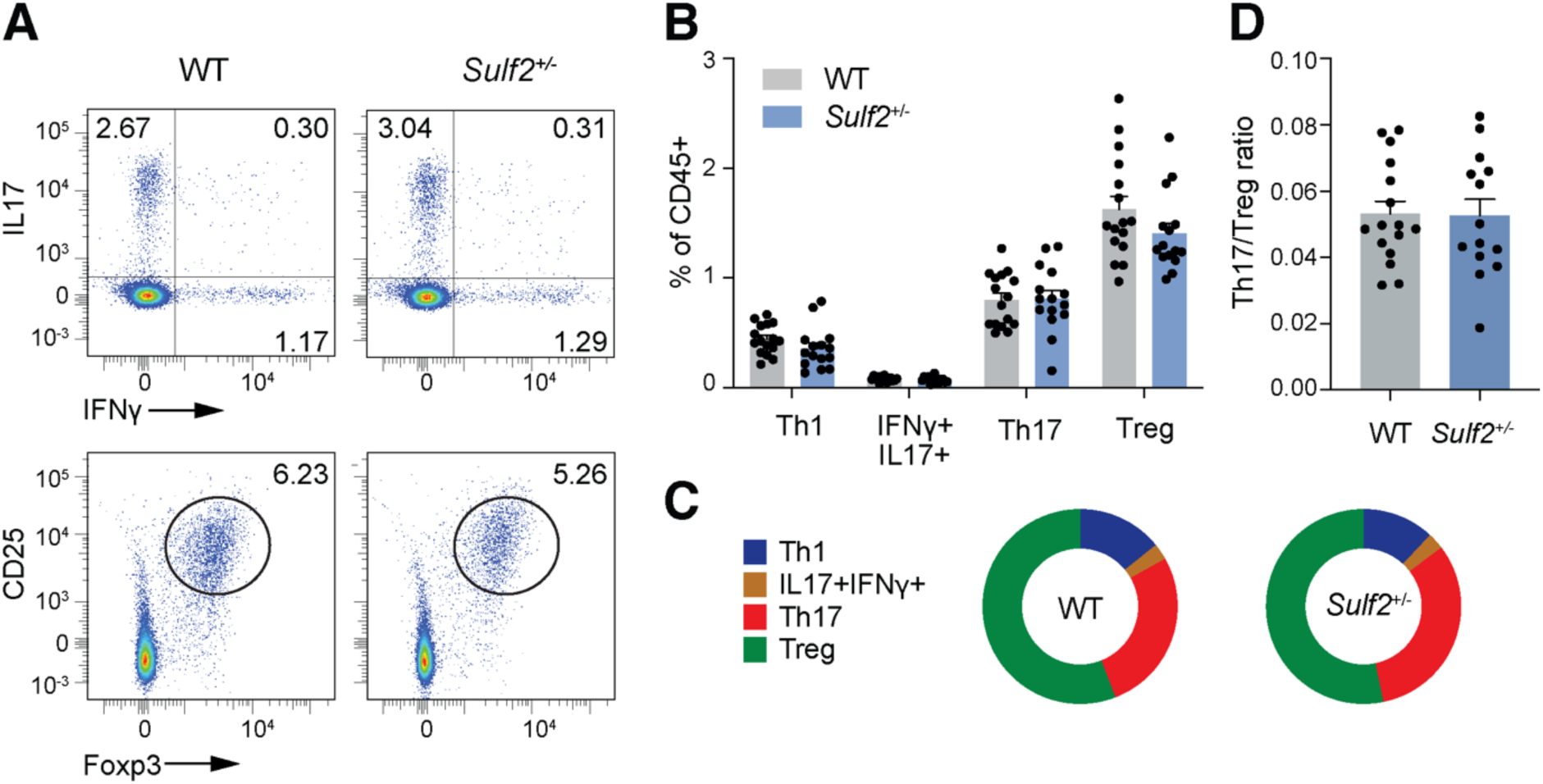
The Th17/Treg ratio in the inguinal lymph nodes of *Sulf2*-deficient mice was unchanged at day 7 of AIA. AIA was induced in WT and *Sulf2^+/-^* bone marrow chimeric mice (n=16). 7 days later, cells were isolated from the inguinal lymph nodes and stimulated for 4 h with PMA (20 ng/ml) and ionomycin (1 mg/ml) in the presence of protein transport inhibitors. CD3+, CD4+ and CD8+ T-cell subsets were analysed by flow cytometry. (**A**). Representative dot plots for Th17 (IFNγ+IL17+) and Treg (CD25+Foxp3+) subsets in inguinal lymph nodes. (**B**). The abundance of T cell subsets (as a percentage of CD45+) was calculated (mean ± SEM, analysed by a two-tailed Student’s *t*-test). (**C**). Graphical representation of data in (**B**). (**D**). The Th17/Treg ratio was calculated from data in (**B**) (mean ± SEM, analysed by two-tailed Student’s *t*-test).

**Supplementary Fig. S8.**
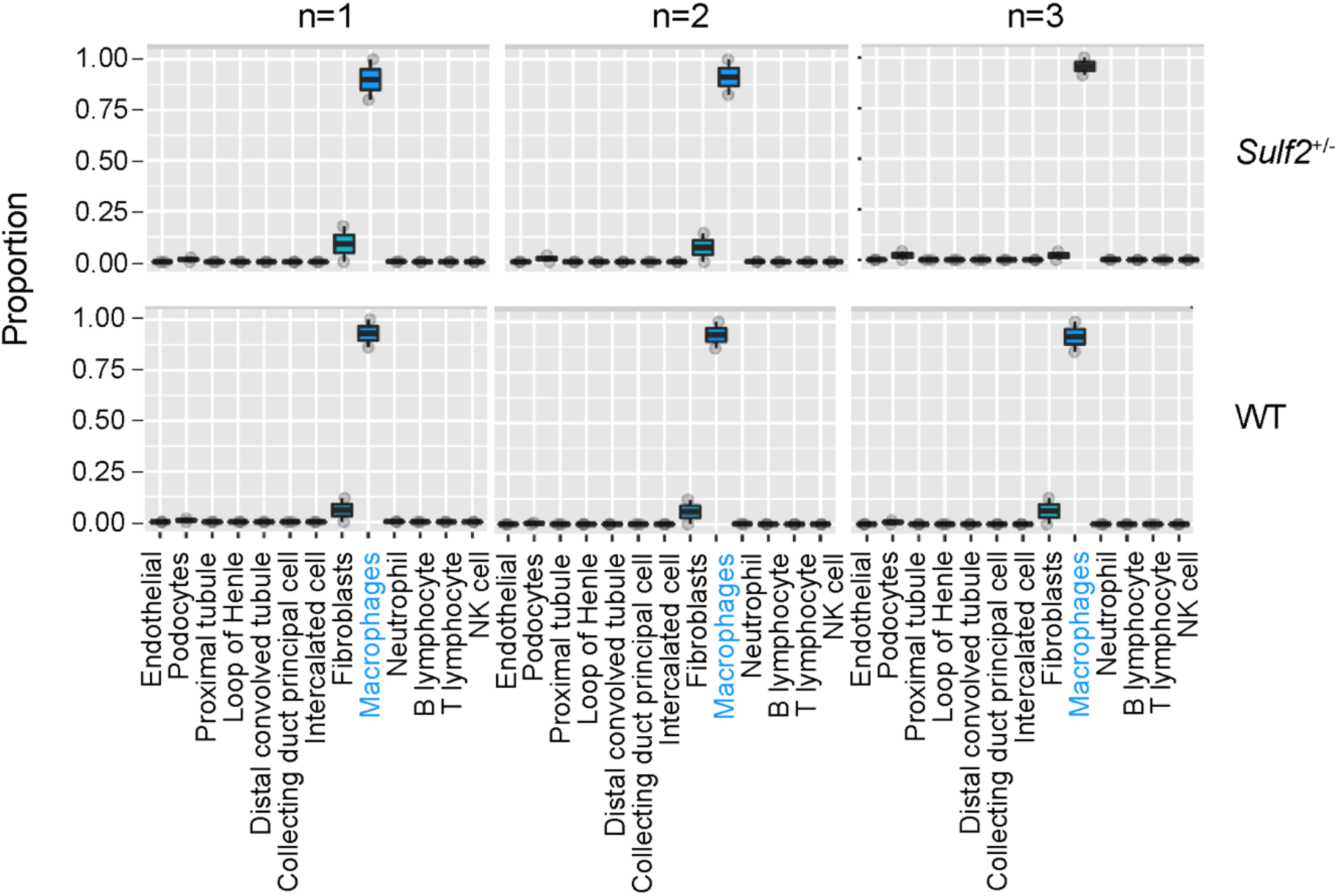
Devolution of bulk RNA sequencing data with MuSIC indicated the majority of transcripts originated from macrophages. The algorithm MuSIC was used to estimate cell type proportions for the bulk RNA sequencing data from *Sulf2*^+/-^ and WT mice (n=3 each) using single cell RNASeq of known cell types [32]. This predicted that the majority (>85%) of cells sequenced were macrophages, with a small proportion of fibroblasts.

**Supplementary Table S1:**
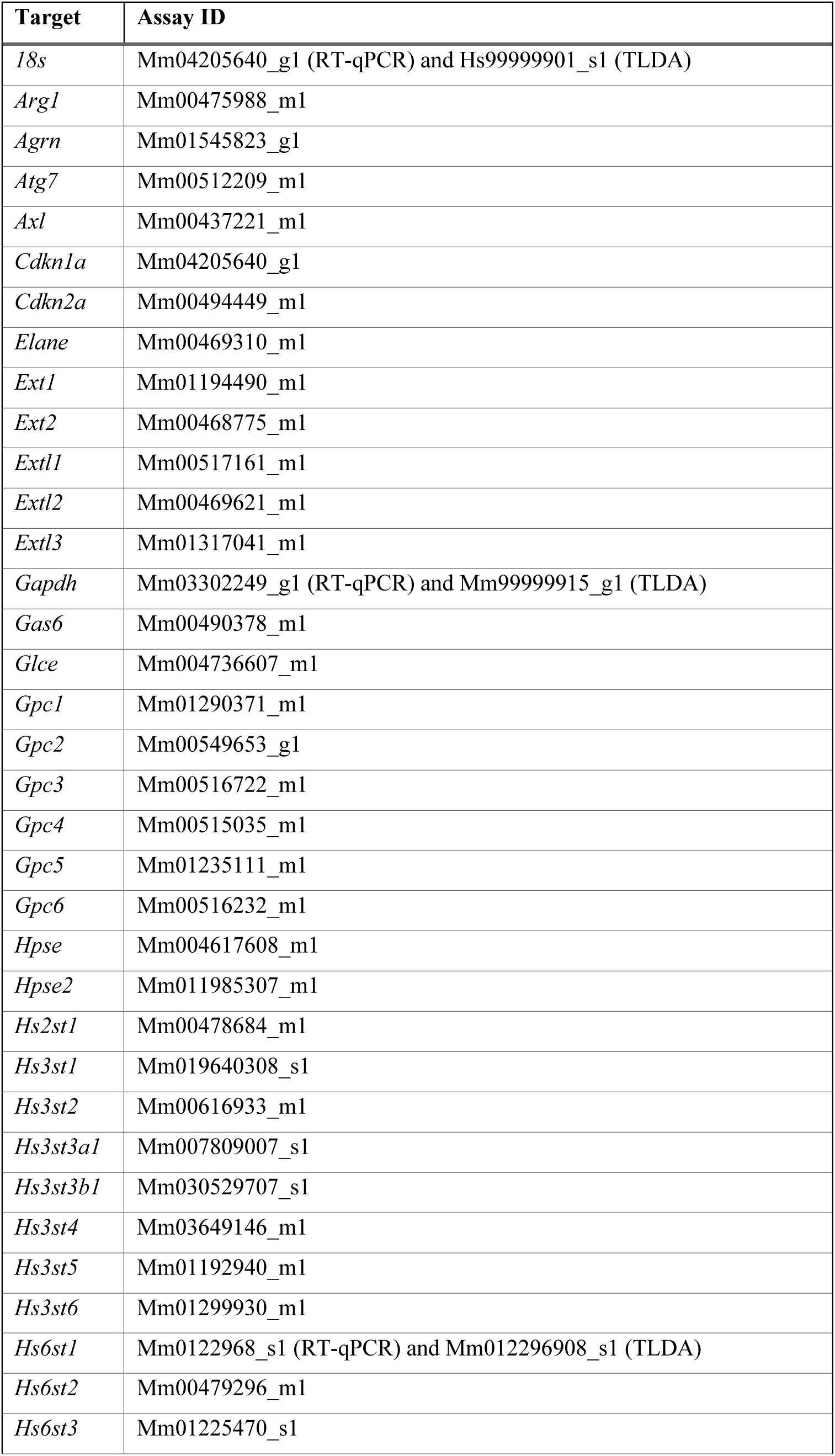
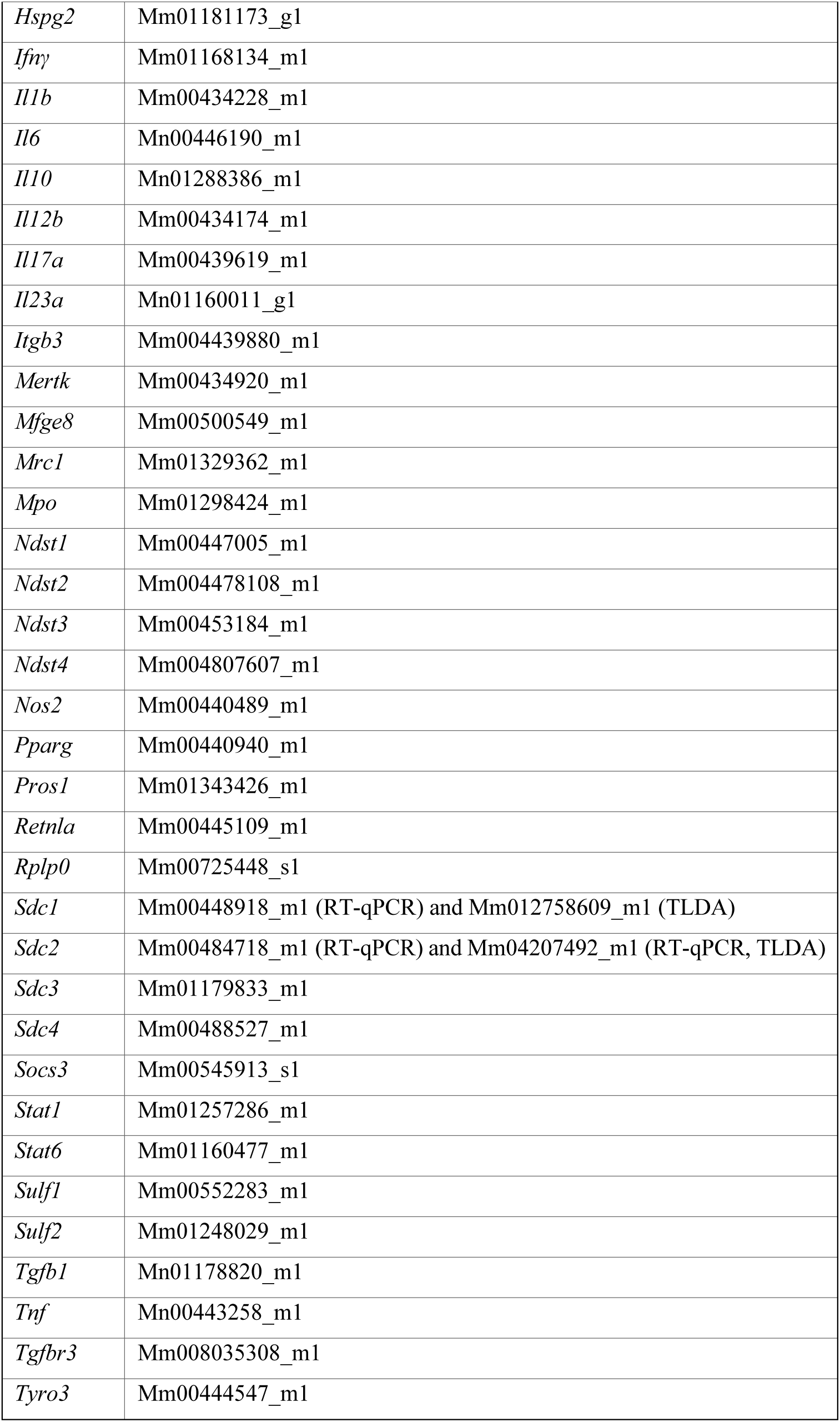
TaqMan primers used to measure gene expression by RT-qPCR and microfluidic cards.

**Supplementary Table S2:**
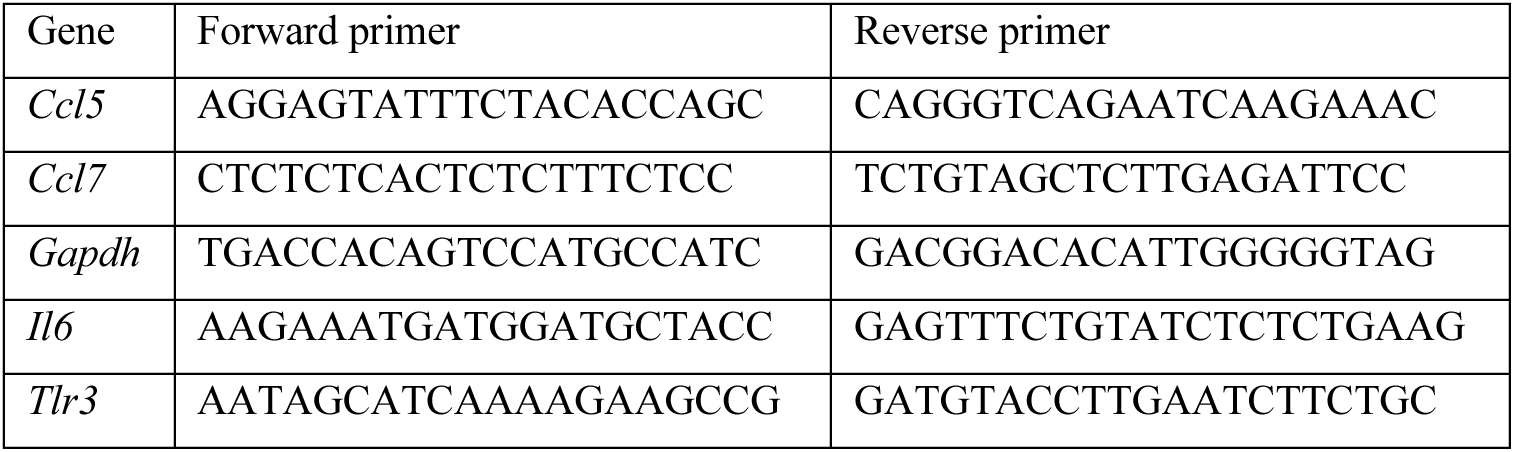
Sybr primers used to measure gene expression by RT-qPCR.

**Supplementary Table S3:**
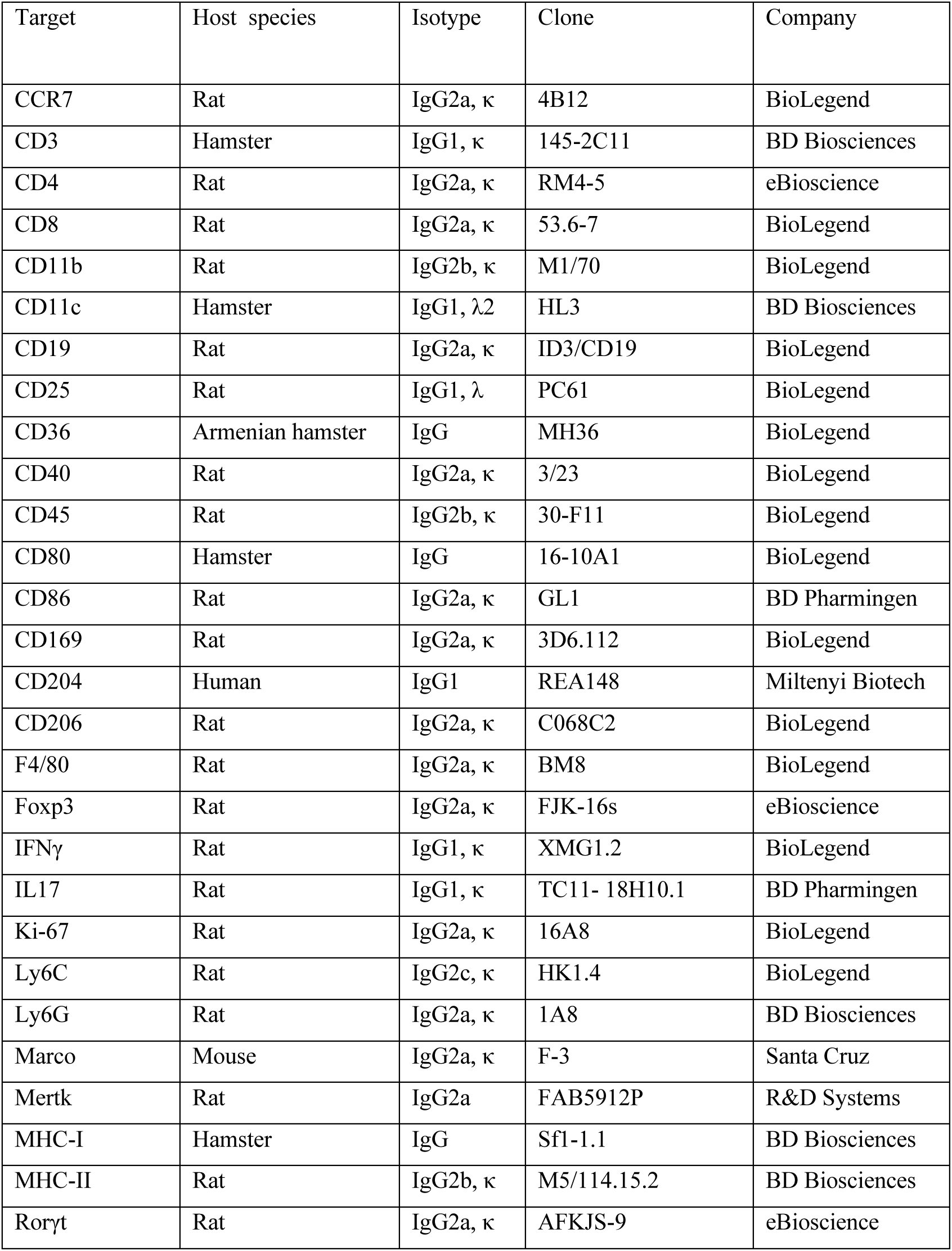
Commercial antibodies used in flow cytometry.

**Supplementary Table S4:** Significantly differentially expressed genes. Bulk RNA sequencing was conducted on cells isolated from knees of WT and *Sulf2*^+/-^ chimeric mice 7 days after initiation of AIA, and differential gene expression analysis performed using DeSeq2.

## References

1. Ori A, Wilkinson MC, Fernig DG (2011) A systems biology approach for the investigation of the heparin/heparan sulfate interactome. J Biol Chem 286: 19892–19904. 10.1074/jbc.M111.228114

2. Collins LE, Troeberg L (2019) Heparan sulfate as a regulator of inflammation and immunity. Journal of Leukocyte Biology. 10.1002/JLB.3RU0618-246R

3. Mulloy B, Rider CC (2006) Cytokines and proteoglycans: An introductory overview. Biochemical Society Transactions 34:409–413. 10.1042/BST0340409

4. Proudfoot A, Johnson Z, Bonvin P, Handel T (2017) Glycosaminoglycan Interactions with Chemokines Add Complexity to a Complex System. Pharmaceuticals 10:70. 10.3390/ph10030070

5. Proudfoot AEI, Handel TM, Johnson Z, et al (2003) Glycosaminoglycan binding and oligomerization are essential for the in vivo activity of certain chemokines. Proc Natl Acad Sci U S A 100:1885–1890. 10.1073/pnas.0334864100

6. Lortat-Jacob H, Baltzer F, Grimaud J-A (1996) Heparin Decreases the Blood Clearance of Interferon-γ and Increases Its Activity by Limiting the Processing of Its Carboxyl-terminal Sequence. J Biol Chem 271:16139–16143. 10.1074/jbc.271.27.16139

7. Kemna J, Gout E, Daniau L, et al (2023) IFNγ binding to extracellular matrix prevents fatal systemic toxicity. Nat Immunol 24:414–422. 10.1038/s41590-023-01420-5

8. Reijmers RM, Groen RWJ, Kuil A, et al (2011) Disruption of heparan sulfate proteoglycan conformation perturbs B-cell maturation and APRIL-mediated plasma cell survival. Blood 117:6162–6171. 10.1182/blood-2010-12-325522

9. 9. Kimberley FC, Van Bostelen L, Cameron K, et al (2009) The proteoglycan (heparan sulfate proteoglycan) binding domain of APRIL serves as a platform for ligand multimerization and cross-linking. FASEB J 23:1584–1595. 10.1096/fj.08-124669

10. Hayashida K, Aquino RS, Park PW (2022) Coreceptor functions of cell surface heparan sulfate proteoglycans. American Journal of Physiology-Cell Physiology 322:C896–C912. 10.1152/ajpcell.00050.2022

11. Li J-P, Kusche-Gullberg M (2016) Heparan Sulfate: Biosynthesis, Structure, and Function. International Review of Cell and Molecular Biology. Elsevier, pp 215–273

12. Poulain FE, Yost HJ (2015) Heparan sulfate proteoglycans: a sugar code for vertebrate development? Development 142:3456–3467. 10.1242/dev.098178

13. Huynh MB, Morin C, Carpentier G, et al (2012) Age-related changes in rat myocardium involve altered capacities of glycosaminoglycans to potentiate growth factor functions and heparan sulfate-altered sulfation. J Biol Chem 287:11363–11373. 10.1074/jbc.M111.335901

14. Huynh MB, Villares J, Sepúlveda Díaz JE, et al (2012) Glycosaminoglycans from aged human hippocampus have altered capacities to regulate trophic factors activities but not Aβ42 peptide toxicity. Neurobiology of Aging 33:1005.e11–1005.e22. 10.1016/j.neurobiolaging.2011.09.030

15. Ghadiali RS, Guimond SE, Turnbull JE, Pisconti A (2017) Dynamic changes in heparan sulfate during muscle differentiation and ageing regulate myoblast cell fate and FGF2 signalling. Matrix Biol 59:54–68. 10.1016/j.matbio.2016.07.007

16. Wijnhoven TJM, Lensen JFM, Rops ALWMM, et al (2006) Aberrant Heparan Sulfate Profile in the Human Diabetic Kidney Offers New Clues for Therapeutic Glycomimetics. American Journal of Kidney Diseases 48:250–261. 10.1053/j.ajkd.2006.05.003

17. Westergren-Thorsson G, Hedström U, Nybom A, et al (2017) Increased deposition of glycosaminoglycans and altered structure of heparan sulfate in idiopathic pulmonary fibrosis. Int J Biochem Cell Biol 83:27–38

18. Tátrai P, Egedi K, Somorácz Á, et al (2010) Quantitative and qualitative alterations of heparan sulfate in fibrogenic liver diseases and hepatocellular cancer. Journal of Histochemistry and Cytochemistry 58:429–441. 10.1369/jhc.2010.955161

19. Hosono-Fukao T, Ohtake-Niimi S, Hoshino H, et al (2012) Heparan sulfate subdomains that are degraded by Sulf accumulate in cerebral amyloid ß plaques of Alzheimer’s disease: evidence from mouse models and patients. Am J Pathol 180:2056–2067. 10.1016/j.ajpath.2012.01.015

20. Chanalaris A, Clarke H, Guimond SES, et al (2019) Heparan Sulfate Proteoglycan Synthesis Is Dysregulated in Human Osteoarthritic Cartilage. American Journal of Pathology 189:632–647. 10.1016/j.ajpath.2018.11.011

21. Ferreras C, Rushton G, Cole CL, et al (2012) Endothelial Heparan Sulfate 6-O-Sulfation Levels Regulate Angiogenic Responses of Endothelial Cells to Fibroblast Growth Factor 2 and Vascular Endothelial Growth Factor. J Biol Chem 287:36132–36146. 10.1074/jbc.M112.384875

22. Swart M, Troeberg L (2019) Effect of Polarization and Chronic Inflammation on Macrophage Expression of Heparan Sulfate Proteoglycans and Biosynthesis Enzymes. Journal of Histochemistry and Cytochemistry. 10.1369/0022155418798770

23. Martinez P, Denys A, Delos M, et al (2015) Macrophage polarization alters the expression and sulfation pattern of glycosaminoglycans. Glycobiology 25:502–513. 10.1093/glycob/cwu137

24. Sikora A-S, Delos M, Martinez P, et al (2016) Regulation of the Expression of Heparan Sulfate 3-O-Sulfotransferase 3B (HS3ST3B) by Inflammatory Stimuli in Human Monocytes. J Cell Biochem 117:1529– 1542. 10.1002/jcb.25444

25. Ng C, Whitelock JM, Williams H, et al (2021) Macrophages bind LDL using heparan sulfate and the perlecan protein core. J Biol Chem 296:100520. 10.1016/j.jbc.2021.100520

26. Gordts PLSM, Foley EM, Lawrence R, et al (2014) Reducing macrophage proteoglycan sulfation increases atherosclerosis and obesity through enhanced type i interferon signaling. Cell Metabolism 20:813–826. 10.1016/j.cmet.2014.09.016

27. Gupta P, Johns SC, Kim SY, et al (2020) Functional Cellular Anti-Tumor Mechanisms are Augmented by Genetic Proteoglycan Targeting. Neoplasia 22:86–97. 10.1016/j.neo.2019.11.003

28. Lum DH, Tan J, Rosen SD, Werb Z (2007) Gene Trap Disruption of the Mouse Heparan Sulfate 6-*O*-Endosulfatase Gene, *Sulf2*. Molecular and Cellular Biology 27:678–688. 10.1128/MCB.01279-06

29. Lamanna WC, Baldwin RJ, Padva M, et al (2006) Heparan sulfate 6-*O*-endosulfatases: discrete *in vivo* activities and functional co-operativity. Biochem J 400:63–73. 10.1042/BJ20060848

30. Reynolds-Peterson CE, Zhao N, Xu J, et al (2017) Heparan sulfate proteoglycans regulate autophagy in *Drosophila*. Autophagy 13:1262–1279. 10.1080/15548627.2017.1304867

31. Morimoto-Tomita M, Uchimura K, Werb Z, et al (2002) Cloning and characterization of two extracellular heparin-degrading endosulfatases in mice and humans. J Biol Chem 277:49175–49185

32. Wang X, Park J, Susztak K, et al (2019) Bulk tissue cell type deconvolution with multi-subject single-cell expression reference. Nat Commun 10:380. 10.1038/s41467-018-08023-x

33. Rusinova I, Forster S, Yu S, et al (2013) Interferome v2.0: an updated database of annotated interferon-regulated genes. Nucleic Acids Res 41:D1040–1046. 10.1093/nar/gks1215

34. You F, Wang P, Yang L, et al (2013) ELF4 is critical for induction of type I interferon and the host antiviral response. Nat Immunol 14:1237–1246. 10.1038/ni.2756

35. Fu XY, Kessler DS, Veals SA, et al (1990) ISGF3, the transcriptional activator induced by interferon alpha, consists of multiple interacting polypeptide chains. Proc Natl Acad Sci USA 87:8555–8559. 10.1073/pnas.87.21.8555

36. El Masri R, Seffouh A, Lortat-Jacob H, Vivès RR (2017) The “in and out” of glucosamine 6-O-sulfation: the 6th sense of heparan sulfate. Glycoconj J 34:285–298

37. Zhang W, Yang F, Zheng Z, et al (2022) Sulfatase 2 Affects Polarization of M2 Macrophages through the IL-8/JAK2/STAT3 Pathway in Bladder Cancer. Cancers 15:131. 10.3390/cancers15010131

38. Holst CR, Bou-Reslan H, Gore BB, et al (2007) Secreted Sulfatases Sulf1 and Sulf2 Have Overlapping yet Essential Roles in Mouse Neonatal Survival. PLoS ONE 2:e575. 10.1371/journal.pone.0000575

39. Ai X, Kitazawa T, Do A-T, et al (2007) SULF1 and SULF2 regulate heparan sulfate-mediated GDNF signaling for esophageal innervation. Development 134:3327–3338. 10.1242/dev.007674

40. HajMohammadi S, Enjyoji K, Princivalle M, et al (2003) Normal levels of anticoagulant heparan sulfate are not essential for normal hemostasis. J Clin Invest 111:989–999. 10.1172/JCI200315809

41. Izvolsky KI, Lu J, Martin G, et al (2008) Systemic inactivation of Hs6st1 in mice is associated with late postnatal mortality without major defects in organogenesis. Genesis 46:8–18. 10.1002/dvg.20355

42. Zec K, Schonfeldova B, Ai Z, et al (2023) Macrophages in the synovial lining niche initiate neutrophil recruitment and articular inflammation. J Exp Med 220:e20220595. 10.1084/jem.20220595

43. Simon J, Surber R, Kleinstäuber G, et al (2001) Systemic Macrophage Activation in Locally-induced Experimental Arthritis. Journal of Autoimmunity 17:127–136. 10.1006/jaut.2001.0534

44. Petrow PK, Thoss K, Katenkamp D, BrÄUer R (1996) Adoptive Transfer of Susceptibility to Antigen-Induced Arthritis into Severe Combined Immunodeficient (SCID) MICE: Role of CD4+ and CD8+ T Cells. LIMM 25:341–353. 10.3109/08820139609059316

45. Guendisch U, Weiss J, Ecoeur F, et al (2017) Pharmacological inhibition of RORγt suppresses the Th17 pathway and alleviates arthritis in vivo. PLoS ONE 12:e0188391. 10.1371/journal.pone.0188391

46. Hashimoto M (2017) Th17 in Animal Models of Rheumatoid Arthritis. JCM 6:73. 10.3390/jcm6070073

47. Korn T, Hiltensperger M (2021) Role of IL-6 in the commitment of T cell subsets. Cytokine 146:155654. 10.1016/j.cyto.2021.155654

48. Tessema M, Yingling CM, Thomas CL, et al (2012) SULF2 methylation is prognostic for lung cancer survival and increases sensitivity to topoisomerase-I inhibitors via induction of ISG15. Oncogene 31:4107– 4116. 10.1038/onc.2011.577

49. Nardelli B, Zaritskaya L, Semenuk M, et al (2002) Regulatory Effect of IFN-K, A Novel Type I IFN, On Cytokine Production by Cells of the Innate Immune System. The Journal of Immunology

50. Axtell RC, Raman C, Steinman L (2013) Type i interferons: Beneficial in Th1 and detrimental in Th17 autoimmunity. Clinical Reviews in Allergy and Immunology 44:114–120. 10.1007/s12016-011-8296-5

51. Saraswat D, Shayya HJ, Polanco JJ, et al (2021) Overcoming the inhibitory microenvironment surrounding oligodendrocyte progenitor cells following experimental demyelination. Nat Commun 12:1923. 10.1038/s41467-021-22263-4

52. Krenn EC, Wille I, Gesslbauer B, et al (2008) Glycanogenomics: A qPCR-approach to investigate biological glycan function. Biochemical and Biophysical Research Communications 375:297–302. 10.1016/j.bbrc.2008.07.144

53. Oshima K, King SI, McMurtry SA, Schmidt EP (2021) Endothelial Heparan Sulfate Proteoglycans in Sepsis: The Role of the Glycocalyx. Semin Thromb Hemost 47:274–282. 10.1055/s-0041-1725064

54. 54. van Det NF, van den Born J, Tamsma JT, et al (1996) Effects of high glucose on the production of heparan sulfate proteoglycan by mesangial and epithelial cells. Kidney Int 49:1079–1089. 10.1038/ki.1996.157

55. Otsuki S, Hanson SR, Miyaki S, et al (2010) Extracellular sulfatases support cartilage homeostasis by regulating BMP and FGF signaling pathways. Proc Natl Acad Sci U S A 107:10202–10207

56. Korf-Klingebiel M, Reboll MR, Grote K, et al (2019) Heparan Sulfate–Editing Extracellular Sulfatases Enhance VEGF Bioavailability for Ischemic Heart Repair. Circ Res 125:787–801. 10.1161/CIRCRESAHA.119.315023

57. Yue X (2017) Epithelial Deletion of Sulf2 Exacerbates Bleomycin-Induced Lung Injury, Inflammation, and Mortality. Am J Respir Cell Mol Biol 57:560–569. 10.1165/rcmb.2016-0367OC

58. Sandoval DR, Gomez Toledo A, Painter CD, et al (2020) Proteomics-based screening of the endothelial heparan sulfate interactome reveals that C-type lectin 14a (CLEC14A) is a heparin-binding protein. J Biol Chem 295:2804–2821. 10.1074/jbc.RA119.011639

59. Tang R, Rosen SD (2009) Functional Consequences of the Subdomain Organization of the Sulfs. J Biol Chem 284:21505–21514. 10.1074/jbc.M109.028472

60. Frese M-A, Milz F, Dick M, et al (2009) Characterization of the Human Sulfatase Sulf1 and Its High Affinity Heparin/Heparan Sulfate Interaction Domain. J Biol Chem 284:28033–28044. 10.1074/jbc.M109.035808

61. El Masri R, Seffouh A, Roelants C, et al (2022) Extracellular endosulfatase Sulf-2 harbors a chondroitin / dermatan sulfate chain that modulates its enzyme activity. Cell Reports. 10.1016/j.celrep.2022.110516

62. Siegel RJ, Singh AK, Panipinto PM, et al (2022) Extracellular sulfatase-2 is overexpressed in rheumatoid arthritis and mediates the TNF-α-induced inflammatory activation of synovial fibroblasts. Cell Mol Immunol 19:1185–1195. 10.1038/s41423-022-00913-x

63. Lallemand Y, Luria V, Haffner-Krausz R, Lonai P (1998) Maternally expressed PGK-Cre transgene as a tool for early and uniform activation of the Cre site-specific recombinase. Transgenic Research 7:105–112. 10.1023/A:1008868325009

64. Topping LM, Romero-Castillo L, Urbonaviciute V, et al (2022) Standardization of Antigen-Emulsion Preparations for the Induction of Autoimmune Disease Models. Front Immunol 13:892251. 10.3389/fimmu.2022.892251

65. Patel H, Ewels P, Peltzer A, et al (2023) nf-core/rnaseq: nf-core/rnaseq v3.11.2 - Resurrected Radium Rhino

66. Park J, Shrestha R, Qiu C, et al (2018) Single-cell transcriptomics of the mouse kidney reveals potential cellular targets of kidney disease. Science 360:758–763. 10.1126/science.aar2131

67. Kanehisa M, Sato Y, Kawashima M (2022) KEGG mapping tools for uncovering hidden features in biological data. Protein Science 31:47–53. 10.1002/pro.4172

68. Heinz S, Benner C, Spann N, et al (2010) Simple combinations of lineage-determining transcription factors prime cis-regulatory elements required for macrophage and B cell identities. Mol Cell 38:576–589. 10.1016/j.molcel.2010.05.004

